# Reciprocal repulsions instruct the precise assembly of parallel hippocampal networks

**DOI:** 10.1101/2020.05.28.122242

**Authors:** Daniel T. Pederick, Jan H. Lui, Ellen C. Gingrich, Chuanyun Xu, Mark J. Wagner, Yuanyuan Liu, Zhigang He, Stephen R. Quake, Liqun Luo

**Affiliations:** Department of Biology, Howard Hughes Medical Institute, Stanford University, Stanford, CA, USA; Neurosciences Graduate Program, Stanford University, Stanford, CA, USA; F.M. Kirby Neurobiology Center, Department of Neurology, Boston Children’s Hospital, Harvard Medical School, Boston, MA, USA; Departments of Bioengineering and Applied Physics, Stanford University, Stanford, CA, USA; Chan Zuckerberg Biohub, Stanford, CA, USA

**Author notes:** Somatosensation and Pain Unit, National Institute of Dental and Craniofacial Research (NIDCR), National Center for Complementary and Integrative Health (NCCIH), National Institutes of Health, Bethesda, MD, USA.

## Abstract

Parallel information processing is a salient feature of complex nervous systems. For example, the medial and lateral hippocampal networks (MHN and LHN) preferentially process spatial- and object-related information, respectively. However, the mechanisms underlying parallel network assembly during development remain largely unknown. Here, we show that complementary expression of cell-surface molecules Teneurin-3 (Ten3) and Latrophilin-2 (Lphn2) in the MHN and LHN, respectively, guides the precise assembly of both the MHN and LHN. Viral-genetic perturbations *in vivo* demonstrate that Ten3+ axons are repelled by target-derived Lphn2, revealing that Lphn2/Ten3-mediated repulsion and Ten3/Ten3-mediated attraction cooperate to control precise target selection of MHN axons. In the LHN, Lphn2+ axons are confined to Lphn2+ targets via repulsion from Ten3+ targets. Our findings demonstrate that assembly of parallel hippocampal networks follows a ‘Ten3→Ten3, Lphn2→Lphn2’ rule instructed by reciprocal repulsions.

## Introduction

Parallel information processing is a salient feature of complex nervous systems, enabling animals to simultaneously process diverse environmental stimuli and efficiently orchestrate appropriate actions. This is achieved through neural networks containing information-processing streams organized in parallel. In order for information to be accurately processed, neural networks must be properly assembled. While remarkable progress has been made in determining how axons are guided to the appropriate anatomical region and select specific target neurons within the region (Kolodkin and Tessier-Lavigne, 2011; Sanes and Zipursky, 2020), these mechanisms have largely been investigated in invertebrate circuits and relatively simple vertebrate circuits owing to their technical ease, with a focus on individual connections. Much less is known about how target selection is achieved in complex networks in the mammalian brain, which contains abundant parallel and often reciprocal connections across multiple regions.

One such parallel organization is present within the hippocampal-entorhinal complex, which is critical for explicit memory and spatial representation (Hafting et al., 2005; O’Keefe and Dostrovsky, 1971; Scoville and Milner, 2000; Squire et al., 2004). Spatial- and object-related information is preferentially processed by two anatomically adjacent parallel networks, the medial and lateral hippocampal networks (MHN and LHN), respectively (Cembrowski et al., 2018; Igarashi et al., 2014). The MHN and LHN consist of topographic projections along the proximal–distal axis between CA1, subiculum, and entorhinal cortex. Specifically, proximal CA1 (pCA1) projects axons to distal subiculum (dSub), while distal CA1 (dCA1) projects axons to proximal subiculum (pSub). Both pCA1 and dSub also form reciprocal connections with the medial entorhinal cortex (MEC), forming the MHN. In parallel, both dCA1 and pSub form reciprocal connections with the lateral entorhinal cortex (LEC), forming the LHN (Naber et al., 2001; Tamamaki and Nojyo, 1995) (Figure 1A).

**Figure 1.**
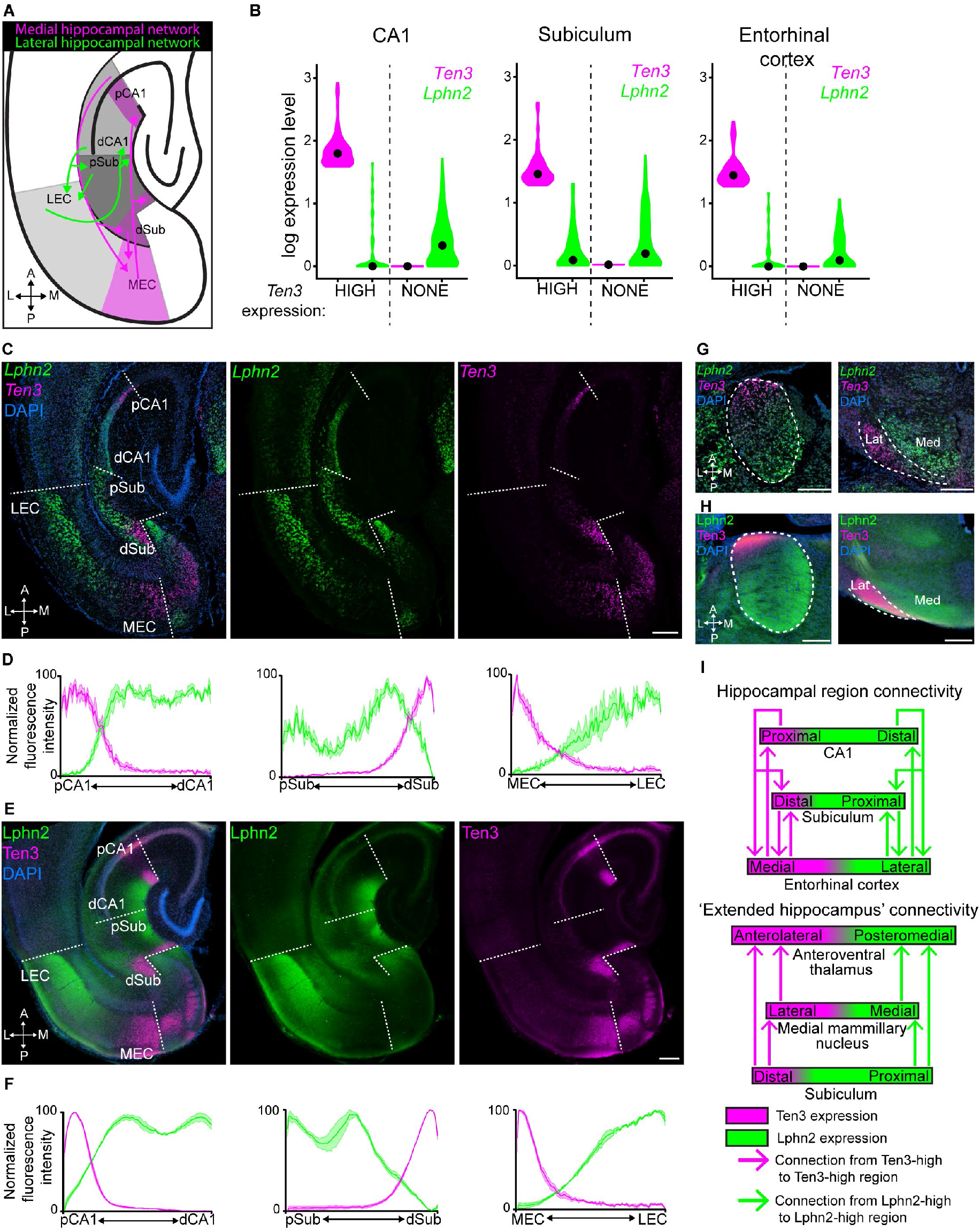
Complementary expression patterns of Lphn2 and Ten3 in the hippocampal network. (**A**) Summary of axonal projection patterns of medial (magenta) and lateral (green) hippocampal network connections. Magenta regions have high Ten3 expression (Berns et al., 2018). A, anterior; P, posterior: L, lateral; M, medial. (**B**) Violin plots highlighting *Lphn2* and *Ten3* expression in *Ten3*-HIGH and *Ten3-* NONE cells across CA1, subiculum, and entorhinal cortex. The unit of expression level is ln [1+ (reads per 10000)]. Black dots represent the median. (**C**) Double *in situ* hybridization for *Lphn2* (middle) and *Ten3* (right) mRNA on a horizontal section of P8 mouse brain. Dashed lines represent boundaries between CA1, subiculum, and entorhinal cortex as labeled in the overlay (left). (**D**) Quantification of *Lphn2* and *Ten3* mRNA across the proximal–distal axis of CA1 (*n* = 3 mice, 29 sections total) and subiculum (*n* = 3 mice, 31 sections total) cell body layers and the medial–lateral axis of layer III entorhinal cortex (*n* = 3 mice, 22 sections total). Mean ± SEM. (**E**) Double immunostaining for Lphn2 (middle; anti-GFP antibody) and Ten3 (right) on a horizontal section of P8 *Lphn2-mVenus* knock-in mouse (Anderson et al., 2017) brain. Dashed lines represent boundaries between CA1, subiculum, and entorhinal cortex as labeled in the overlay (left). (**F**) Quantification of Lphn2 and Ten3 protein across the proximal–distal axis of molecular layers of CA1 (*n* = 3 mice, 23 sections total) and subiculum (*n* = 3 mice, 22 sections total), and the medial– lateral axis of layer III entorhinal cortex (*n* = 3 mice, 14 sections total). Mean ± SEM. (**G**) Double *in situ* hybridization showing *Lphn2* and *Ten3* mRNA expression in the anteroventral thalamus (left) and medial mammillary nucleus (right) of a P8 mouse. Lat: lateral; Med; medial. (**H**) Double immunostaining for Lphn2 (left; anti-GFP antibody) and Ten3 on a P8 *Lphn2-mVenus* knock-in mouse (Anderson et al., 2017) brain, showing protein expression in anteroventral thalamus (left) and the medial mammillary nucleus (right). Dashed lines represent anteroventral nucleus (left) and the boundary between the medial and lateral regions of the medial mammillary nucleus (right) in both (**G**) and (**H**). (**I**) Schematic summary of the expression pattern of Lphn2 and Ten3 in relation to interconnected regions of the lateral and medial hippocampal networks. Scale bars, 200 μm.

We have previously shown that the type II transmembrane protein Tenuerin-3 (Ten3) has matching expression in all interconnected regions of the MHN (Figure S1A). Ten3 is required in both pCA1 and dSub for the precise target selection of the pCA1→dSub axons, and promotes aggregation of non-adhesive cells (Berns et al., 2018). These data support a homophilic attraction mechanism by which Ten3 regulates the pCA1→dSub axon projection in the precise assembly of the MHN. It remains unclear whether matching gene expression exists in the LHN and how this contributes to parallel hippocampal network assembly. Does the assembly of the LHN utilize a mechanism similar to or distinct from that of the MHN? Are the assemblies of MHN and LHN coordinated?

## Results

### Complementary Lphn2/Ten3 expression across parallel hippocampal networks

We hypothesized that cell-surface molecules that are preferentially expressed in the LHN and thus have inverse expression to Ten3 may play a role in the precise assembly of parallel hippocampal networks. To identify such molecules, we performed fluorescence-activated cell sorting-based single-cell RNA sequencing (scRNA-seq) of postnatal day 8 (P8) excitatory neurons, in MHN subregions (pCA1, dSub, and MEC) or LHN subregions (dCA1, pSub, and LEC) (see figs. S1, S2, and table S1 for details). We used the ‘ground-truth’ that Ten3 has enriched expression in all MHN subregions (Berns et al., 2018) to specifically search for genes with inverse expression patterns to *Ten3* in CA1, subiculum, and entorhinal cortex. We identified *Lphn2*, an adhesion G-protein-coupled receptor (GPCR) known to bind Teneurins (Boucard et al., 2014; Li et al., 2018; O’Sullivan et al., 2012; Sando et al., 2019; Silva et al., 2011; del Toro et al., 2020), as one of the few cell-surface molecules that showed inverse expression to *Ten3* in CA1, subiculum, and entorhinal cortex (Figure 1B and table S1). Teneurins are known to interact in *trans* with Latrophilins, and this interaction has been implicated in multiple neurodevelopmental processes (Sando et al., 2019; del Toro et al., 2020; Vysokov et al., 2018). This finding suggests that Lphn2 may participate in guiding the assembly of hippocampal networks.

To validate the finding from our scRNA-seq screen, we performed *in situ* expression analysis on P8 brains. Double *in situ* hybridization for *Lphn2* and *Ten3* mRNA revealed specific expression of *Lphn2* in dCA1, pSub, and LEC, which was complementary to *Ten3* enrichment in pCA1, dSub, and MEC (Figure 1C and Figure S3A). Quantification of *Lphn2* and *Ten3* mRNA showed opposing gradients of expression, with *Lphn2* expression levels increasing more sharply after the *Ten3*-high zone in CA1 and subiculum compared to entorhinal cortex (Figure 1D and Figure S3B). We also examined protein expression using an anti-Ten3 antibody (Berns et al., 2018) and an anti-GFP antibody in *Lphn2-mVenus-*knockin mice (Anderson et al., 2017). In all regions, Lphn2 and Ten3 proteins were expressed in the synaptic layers corresponding to their mRNA expression, including molecular layer of CA1, cell body and molecular layers of the subiculum, and layer III of the entorhinal cortex (Figure 1E and F and Figure S3C and D).

The hippocampal networks include extended projections from the subiculum to the anteroventral thalamus and the medial mammillary nucleus (Ishizuka, 2001; Meibach and Siegel, 1977; Witter and Groenewegen, 1990; Wright et al., 2010, 2013). These projections also display targeting specificity within the medial and lateral hippocampal networks and can be highlighted by tracing from the *Lphn2+* and *Ten3+* regions of the subiculum (Figure S4). Double *in situ* hybridization for *Lphn2* and *Ten3* mRNA in subiculum target regions of P8 mice revealed that *Lphn2* and *Ten3* expression matched axonal projections from neurons in the *Lphn2+* and *Ten3+* subiculum regions. *Lphn2* showed enriched expression in the posteromedial portion of the anteroventral thalamus and the medial part of the medial mammillary nucleus, coinciding with axon projection targets of proximal, *Lphn2+* subiculum neurons. By contrast, *Ten3* was highly expressed in the anterolateral portion of the anteroventral thalamus and the lateral part of the medial mammillary nucleus, coinciding with axon projection targets of distal, *Ten3+* subiculum neurons (compare Figure 1G with Figure S4C and D). Expression of Lphn2 and Ten3 protein was consistent with mRNA expression in extended hippocampus target regions (Figure 1G and H).

In summary, *Lphn2* and *Ten3* mRNA, as well as Lphn2 and Ten3 proteins, exhibit striking complementary expression in multiple regions of the developing extended hippocampal networks, including CA1, subiculum, entorhinal cortex, anteroventral thalamus, and medial mammillary nucleus. In all cases, the connection specificity follows a ‘Ten3→Ten3, Lphn2→Lphn2’ rule that correlates cell surface molecule expression with connectivity (Figure 1I).

### Subiculum Lphn2 repels Ten3+ CA1 axons

Ten3 directs targeting of pCA1→dSub axons in the MHN through matching expression and homophilic attraction (Berns et al., 2018). Does Lphn2 also mediate homophilic attraction to assemble the LHN? To test this, we performed an *in vitro* cell aggregation assay using non-adhesive K562 cells. We found that Ten3-expressing K562 cells formed aggregates as previously reported (Berns et al., 2018), but Lphn2-expressing cells did not (Figure S5A and B). However, Ten3-expressing cells aggregated with Lphn2-expressing cells (Figure S5A and B), consistent with the previously reported heterophilic interaction between Teneurins and Latrophilins (Anderson et al., 2017; Berns et al., 2018; Boucard et al., 2014; Li et al., 2018; del Toro et al., 2020). The heterophilic interaction of Ten3 and Lphn2 combined with their complementary expression in the MHN versus LHN suggests that the interaction between Lphn2 and Ten3 may result in repulsion, which could allow distinct target selection of the MHN and LHN.

To determine if target-derived Lphn2 repels Ten3+ axons, we interrogated the CA1→Sub connection. Ten3+ pCA1 and Lphn2+ mid-CA1 (mCA1) axons extend along a tract above the subiculum cell body layer until they reach Ten3+ dSub and Lphn2+ mid-subiculum (mSub), respectively, where they invade the cell body layer of the subiculum to form synapses (Berns et al., 2018) (Figure 2A). Because the CA1→Sub projection develops postnatally (Berns et al., 2018), we injected lentivirus (LV) expressing *GFP* (control) or *GFP-P2A-Lphn2* into the Lphn2-low dSub of mice at P0 to create a region of subiculum expressing Lphn2 across the entire proximal–distal axis. We then injected adeno-associated virus expressing membrane-bound mCherry (*AAV-ChR2-mCh*) into pCA1 in these same mice as adults to label and trace Ten3+ CA1 axons (Figure 2B). The portion of subiculum transduced by LV was only a subset of the total pCA1 axon targeting region along the orthogonal medial–lateral axis, allowing us to determine if pCA1 axons target LV-transduced subiculum regions differently compared to neighboring non-LV-transduced subiculum regions. In *LV-GFP* controls, pCA1 axon targeting was unaffected by GFP expression (Figure 2C and Figure S6A and B). By contrast, in *LV-GFP-P2A-Lphn2* injected brains, pCA1 axons avoided regions where Lphn2 was misexpressed in dSub (Figure 2E and Figure S6D, E).

**Figure 2.**
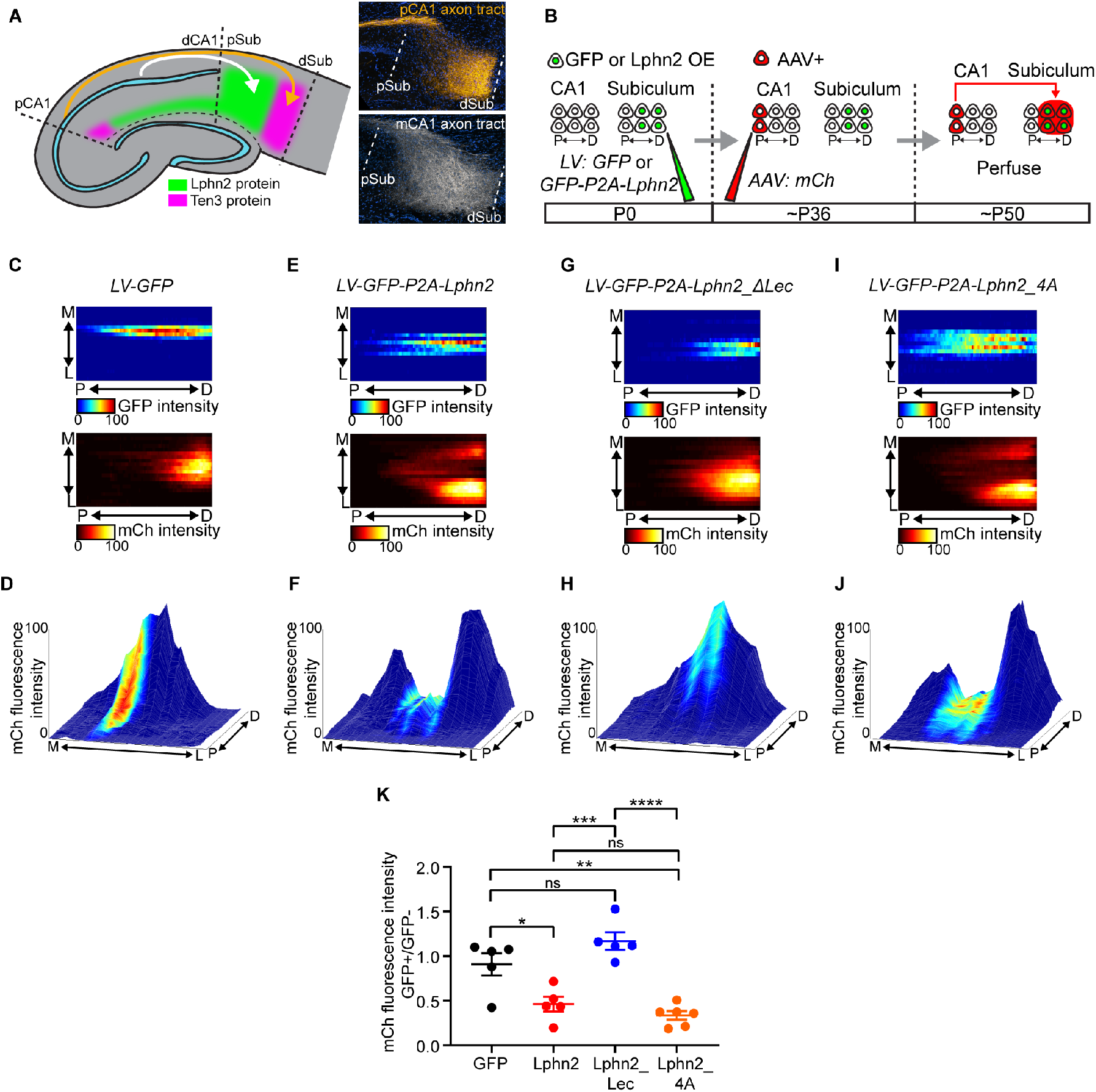
Ten3+ pCA1 axons avoid distal subiculum misexpressing Lphn2 in a Lphn2/Teneurin interaction–dependent and Lphn2/FLRT interaction–independent manner. (**A**) Left, schematic highlighting the trajectory of pCA1 (orange) and mCA1 (white) axons to the subiculum. Right, confocal images of adult subiculum showing pCA1 axons (orange, top) and mCA1 (white, bottom) extending along a tract above the subiculum cell body layer until they turn into dSub and mSub target areas, respectively. (**B**) Experimental design. (**C**), (**E**), (**G**), (**I**) Heatmaps showing normalized GFP fluorescence intensity (top) and normalized mCh fluorescence intensity (bottom) in subiculum. Each row is one section; adjacent sections are separated by 120 μm. Lentiviral (LV) constructs injected are above each pair of panels. P, proximal; D, distal; M, medial; L, lateral. (**D**), (**F**), (**H**), (**J**) Mountain-plots showing normalized GFP fluorescence intensity as color and normalized mCh fluorescence intensity as height. Same data as (**C**), (**E**), (**G**), (**I**), respectively. (**K**) Ratio of mCh fluorescent intensity of GFP+ versus GFP– regions. *LV-GFP* (n = 5), *LV-GFP-P2A-Lphn2* (n = 5), *LV-GFP-P2A-Lphn2_ΔLec* (n = 5) and *LV-GFP-P2A-Lphn2_4A* (n = 6). Mean ± SEM; one-way ANOVA with Tukey’s multiple comparisons test. \*\*\*\**P* ≤ 0.0001; *** *P* ≤ 0.001; ** *P* ≤ 0.01; * *P* ≤ 0.05; ns, not significant.

To further assess how Ten3+ axons target the subiculum in regions where Lphn2 is misexpressed, we generated ‘mountain-plots,’ where pCA1 axons (mCh) and LV injection site (GFP) intensities were plotted on the same graph as height and color, respectively. Expression of GFP alone did not affect the intensity of pCA1 axons targeting the subiculum (Figure 2D and Figure S6C). However, pCA1 axon intensity was markedly reduced in dSub regions misexpressing Lphn2 (Figure 2F and Figure S6F; quantified in Figure 2K). These data suggest that Ten3+ axons are repelled by misexpressed Lphn2 at the dSub target.

### Repulsion requires Lphn2/Teneurin but not Lphn2/FLRT interaction

To test whether Lphn2-mediated repulsion requires Lphn2/Ten3 interaction, we utilized a deletion of the lectin binding domain in Latrophilins, which has been shown to abolish Teneurin binding without affecting interactions with other known partners or surface expression (Boucard et al., 2014; Li et al., 2018; Sando et al., 2019; del Toro et al., 2020). We validated in our K562 cell aggregation assay that Lphn2_ΔLec disrupted Ten3 interaction without affecting interaction with FLRT2 (Figure S5C and D), a member of the fibronectin leucine-rich transmembrane protein family known to bind Latrophilins (Jackson et al., 2015; O’Sullivan et al., 2012). We then misexpressed Lphn2_ΔLec in subiculum to determine if pCA1 axon avoidance depends on a Lphn2/Teneurin interaction. We found that in *LV-GFP-P2A-Lphn2_ΔLec* injected brains, Ten3+ pCA1 axons were not affected by Lphn2_ΔLec expression in dSub and displayed normal targeting to dSub (Figure 2G and Figure S7A and B). Mountain plot analysis also indicated that misexpressed Lphn2_ΔLec did not repel Ten3+ pCA1 axons (Figure 2H and K; Figure S7C).

Recently, FLRTs have been shown to coincidently interact with Teneurin and Latrophilin to direct synapse specificity and repulsive guidance for migrating neurons (Del Toro et al., 2020; Sando et al., 2019). Intriguingly, expression of *FLRT2* was enriched in Ten3-high CA1 cells (Figure S8A), suggesting that it may play a role in pCA1 axon repulsion from Lphn2. Mutation of four residues in the olfactomedin domain of Latrophilin to alanines has been show to abolish FLRT-Lphn binding while maintaining Teneurin binding and surface expression (Lu et al., 2015). We validated that in K562 cells, Lphn2_4A disrupted FLRT2 binding without affecting Ten3 binding (Figure S5E and F). Yet, overexpression of Lphn2_4A in the subiculum still caused a decrease of pCA1 axon intensity in GFP+ dSub regions compared to adjacent GFP–dSub regions (Figure 2I and Figure S7D and E) to the same extent as wild-type Lphn2 (Figure 2K). These gain-of-function experiments suggest that repulsion of Ten3+ pCA1 axons by target-derived Lphn2 requires Lphn2/Teneurin but not Lphn2/FLRT interaction.

### Ten3+ CA1 axons invade *Lphn2-null* subiculum targets

To determine if endogenous *Lphn2* in the subiculum target is necessary for correct pCA1→dSub targeting, we performed a loss-of-function experiment by injecting *LV-GFP-Cre* into the subiculum of control and *Lphn2^fl/fl^* mice (Anderson et al., 2017) at P0, followed by *AAV-ChR2-mCh* in pCA1 of the same mice as adults to assess Ten3+ axon targeting (Figure 3A). In *Lphn2^+/+^* control mice, pCA1 axons targeted dSub and were not disrupted when projecting into GFP-Cre+ expressing regions (Figure 3B). By contrast, Ten3+ pCA1 axons spread into the pSub of GFP-Cre+ regions in *Lphn2^fl/fl^* mice (Figure 3C). Quantification of pCA1 axon intensity in GFP-Cre+ sections revealed that pCA1 axons in *Lphn2^fl/fl^* mice had increased intensity in the more proximal regions and decreased intensity in the most distal region of the subiculum compared to *Lphn2^+/+^* mice (Figure 3E and F; red vs black). These data suggest that Lphn2 in pSub normally repels Ten3+ pCA1 axons, enabling them to specifically target dSub.

**Figure 3.**
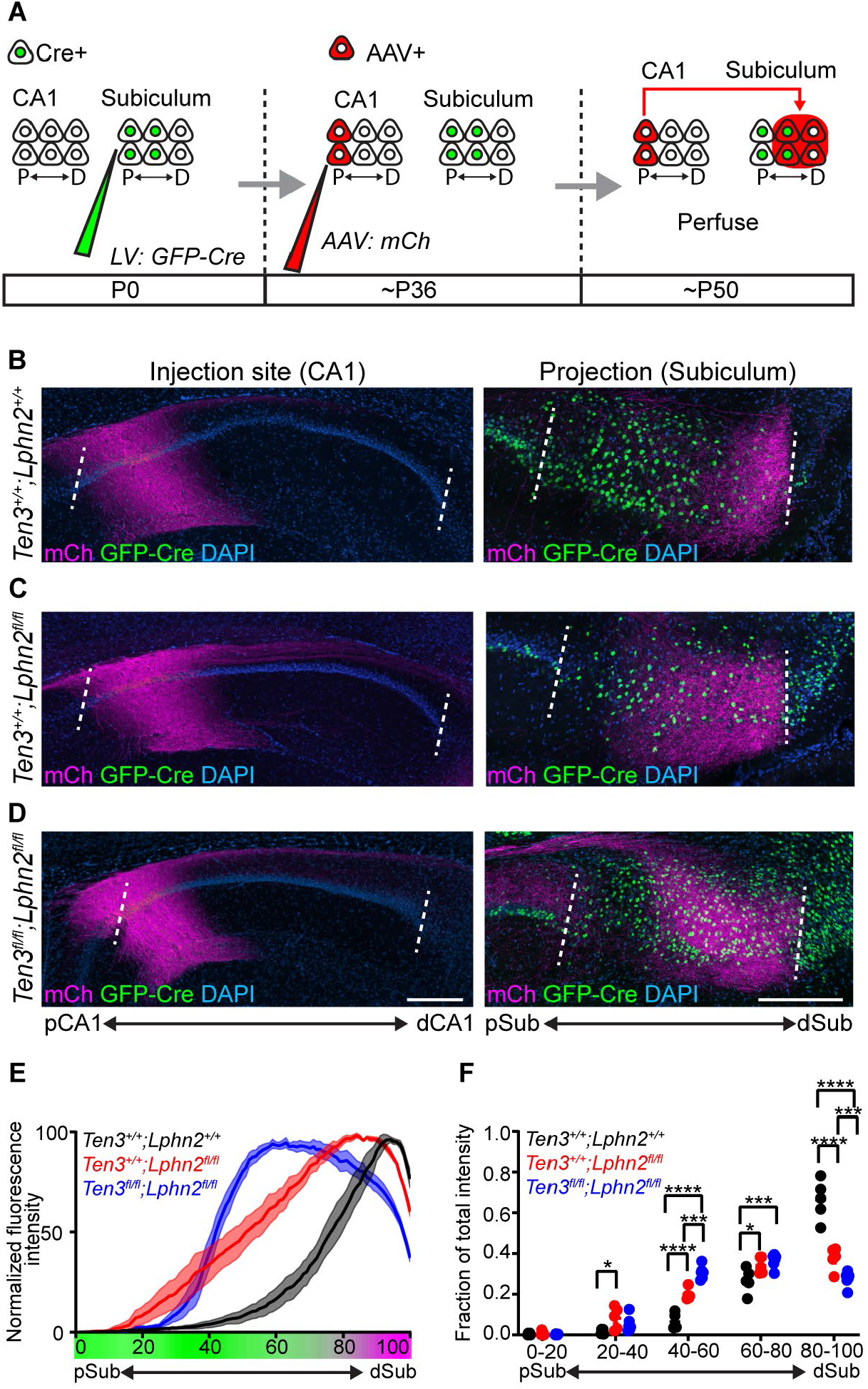
Lphn2/Ten3-mediated repulsion and Ten3/Ten3-mediated attraction cooperate to guide pCA1→dSub target selection. (**A**) Experimental design. (**B** to **D**) Representative images of *AAV-mCh* (magenta) injections in pCA1 (left) and corresponding projections overlapping with *LV-GFP-Cre* (green) injection sites in the subiculum (right). Genetic conditions are as indicated. (**E**) Normalized mean fluorescence intensity traces of subiculum projections from pCA1 in GFP-Cre+ sections for *Lphn2^+/+^* (n = 5 mice), *Lphn2^fl/fl^* (n = 5 mice) and *Lphn2^fl/fl^;Ten3^fl/fl^* (n = 6 mice). Mean ± SEM. Color bar under x-axis represents Lphn2 (green) and Ten3 (magenta) expression in subiculum as quantified in Figure S3. (**F**) Fraction of total axon intensity for the same data as (**E**) across 20 percent intervals. Mean ± SEM, two-way ANOVA with Sidak’s multiple comparisons test. \*\*\*\**P* ≤ 0.0001; *** *P* ≤ 0.001; * *P* ≤ 0.05. Scale bar, 200 μm. Injection site locations in CA1 are shown in Figure S1

To rule out the possibility that the ectopic invasion of pCA1 axons into *Lphn2^−/−^* pSub results from loss of Lphn2 interaction with a molecule other than Ten3 [e.g., another Teneurin that is expressed in pCA1 (Figure S8B)], we performed the same *Lphn2* loss-of-function experiment in *Ten3^−/−^* mice. Anterograde tracing from pCA1 in *Lphn2^+/+^;Ten3^−/−^* mice showed that pCA1 axons spread more along the proximal–distal axis of the subiculum as previously reported (Berns et al., 2018) (Figure S9A). In *Lphn2^fl/fl^; Ten3^−/−^* mice, pCA1 axons also showed similar spreading (Figure S9B). We observed no significant increase of axon intensity in the proximal subiculum of *Lphn2^fl/fl^;Ten3^−/−^* mice compared with *Lphn2^+/+^;Ten3^−/−^* mice (Figure S9C and D). The lack of an additional axon mistargeting phenotype in *Lphn2^fl/fl^; Ten3^−/−^* mice compared to *Lphn2^+/+^;Ten3^−/−^* mice suggests that Ten3 is required for the effect of loss of subiculum Lphn2 on pCA1 axon targeting, and that Lphn2/Ten3-mediated repulsion instructs precise pCA1→dSub target selection.

### Lphn2/Ten3-mediated repulsion and Ten3/Ten3-mediated attraction cooperate

Loss of Lphn2/Ten3 heterophilic repulsion (above) or Ten3 homophilic attraction (Berns et al., 2018) alone both disrupt precise pCA1→dSub axon targeting. What is the relative contribution of each? To test this, we simultaneously conditionally deleted both *Lphn2* and *Ten3* in the subiculum and assessed the targeting of Ten3+ pCA1 axons. We found that pCA1 axons projecting into GFP-Cre+ regions of *Lphn2^fl/fl^;Ten3^fl/fl^* mice targeted more proximal regions of subiculum and also had decreased fluorescence intensity in dSub (Figure 3D). Quantification of pCA1 axons in GFP-Cre+ subiculum sections of *Lphn2^fl/fl^;Ten3^fl/fl^* mice showed a significant increase in axon intensity into the *Lphn2^−/−^* pSub region compared to *Lphn2^+/+^;Ten3^+/+^* mice (Figure 3E and F; blue vs black) and additionally had decreased fluorescence intensity in the *Ten3^−/−^* dSub region compared to *Lphn2^fl/fl^;Ten3^+/+^* mice (Figure 3E and F; blue vs red).

The increase of pCA1 axon intensity in the pSub of *Lphn2^fl/fl^;Ten3^fl/fl^* compared to *Lphn2^+/+^;Ten3^+/+^* confirms the loss of repulsion of pCA1 axons from Lphn2+ pSub. At the same time, the decrease of axon intensity in dSub of *Lphn2^fl/fl^;Ten3^fl/fl^* compared to *Lphn2^fl/fl^;Ten3^+/+^* indicates a loss of attraction of Ten3+ pCA1 axons to Ten3+ dSub. Thus, Lphn2/Ten3-mediated heterophilic repulsion and Ten3/Ten3-mediated homophilic attraction cooperate in orchestrating the precise targeting of pCA1 axons to dSub.

### Subiculum Ten3 repels Lphn2+ CA1 axons

In addition to serving as a repulsive ligand for target selection of Ten3+ MHN neurons, could Lphn2 also act as a receptor to regulate target selection of LHN neurons? Specifically, could Lphn2+ axons be repelled from Ten3+ targets to regulate the precision of LHN connections? To test this, we injected *LV-GFP-Cre* into the subiculum of *Ten3^+/+^* (control) and *Ten3^fl/fl^*mice at P0, followed by *AAV-ChR2-mCh* in mCA1 of the same mice as adults to assess Lphn2+ mCA1 axon targeting (Figure 4A). In *Ten3^+/+^* mice, mCA1 axons predominantly projected to mSub (Figure 4B). However, in *Ten3^fl/fl^* mice, mCA1 axons spread into *Ten3*-null dSub (Figure 4C). Quantification of axons in the subiculum showed a significant increase in axon intensity in the dSub of *Ten3^fl/fl^* mice compared to *Ten3^+/+^*mice (Figure 4D and E). Thus, Ten3 in dSub prevents Lphn2+ mCA1 axon invasion into dSub.

**Figure 4.**
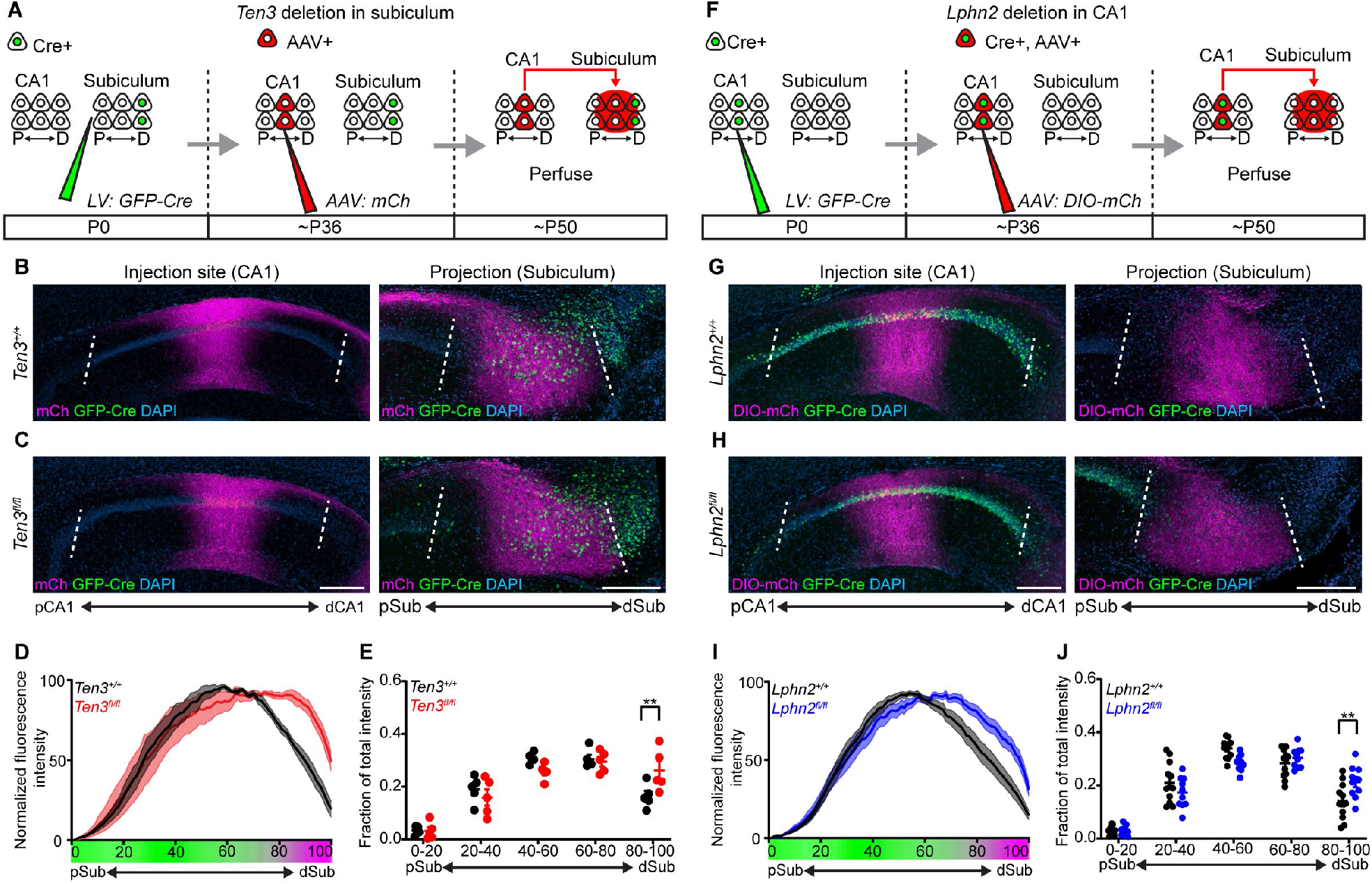
Lphn2+ mCA1 axons avoid Ten3+ dSub, contributing to LHN target selection. (**A**) Experimental design for deleting *Ten3* in the subiculum and tracing mCA1 axons. (**B** and **C**) Representative images of *AAV-mCh* (magenta) injections in mCA1 (left) and corresponding projections overlapping with *LV-GFP-Cre* (green) injection sites in the subiculum (right). Genetic conditions are as indicated. (**D**) Normalized mean fluorescence intensity traces of subiculum projections from mCA1 in GFP-Cre+ sections for *Ten3^+/+^* (n = 5 mice) and *Ten3^fl/fl^* (n = 5 mice). Mean ± SEM. Color bar under x-axis represents Lphn2 (green) and Ten3 (magenta) expression in subiculum as quantified in Figure S3. (**E**) Fraction of total axon intensity for the same data as (**D**) across 20 percent intervals. Mean ± SEM; two-way ANOVA with Sidak’s multiple comparisons test, ** *P* ≤ 0.01. (**F**) Experimental design for deleting *Lphn2* in CA1 and tracing *Lphn2*-null mCA1 axons. (**G** and **H**) Representative images of *AAV-DIO-mCh* (magenta) injections in mCA1 (left) and corresponding projections in the subiculum (right). Genetic conditions are as indicated. (**I**) Normalized mean fluorescence intensity traces of subiculum projections from mCA1 axons for *Lphn2^+/+^* (n = 12 mice) and *Lphn2^fl/fl^* (n = 10 mice). Mean ± SEM. Color bar under x-axis represents Lphn2 (green) and Ten3 (magenta) expression in subiculum as quantified in Figure S3. (**J**) Fraction of total axon intensity for the same data as (**I**) across 20 percent intervals. Mean ± SEM; two-way ANOVA with Sidak’s multiple comparisons test; ** *P* ≤ 0.01. Scale bars, 200 μm. Injection site locations in CA1 are shown in Figure S10.

To test if Lphn2 in mCA1 axons is required for their target precision, we deleted *Lphn2* from CA1 followed by tracing of *Lphn2-null* mCA1 axons (Figure 4F). Control mCA1 axons targeted mSub (Figure 4G), whereas *Lphn2*-null mCA1 axons spread into the most distal subiculum (Figure 4H, quantified in Figure 4I and J). Thus, Lphn2 is cell-autonomously required in mCA1 neurons to prevent their axons from invading Ten3+ dSub. Taken together with the *Ten3* conditional deletion in subiculum above, these data indicate that Lphn2+ mCA1 axons are repelled by target-derived Ten3.

## Discussion

Here, we provide multiple lines of evidence demonstrating that parallel hippocampal networks are assembled through reciprocal repulsions between Ten3- and Lphn2-expressing cells. Our results demonstrate that Lphn2 and Ten3 instruct the assembly of both MHN and LHN connections from CA1 to the subiculum (Figure 5A). Specifically, in the MHN, Lphn2/Ten3-mediated heterophilic repulsion and Ten3/Ten3-mediated homophilic attraction cooperate to instruct pCA1→dSub targeting; in the LHN, Ten3/Lphn2-mediated heterophilic repulsion confines Lphn2+ axons to the Lphn2+ target region, contributing to the precision of mCA1→mSub targeting (Figure 5B). Together, these data show that the mechanisms required for parallel network assembly in the hippocampus are intertwined, utilizing multiple interactions of two cell-surface molecules and reciprocal repulsions to ensure precise segregated connections are formed.

**Figure 5.**
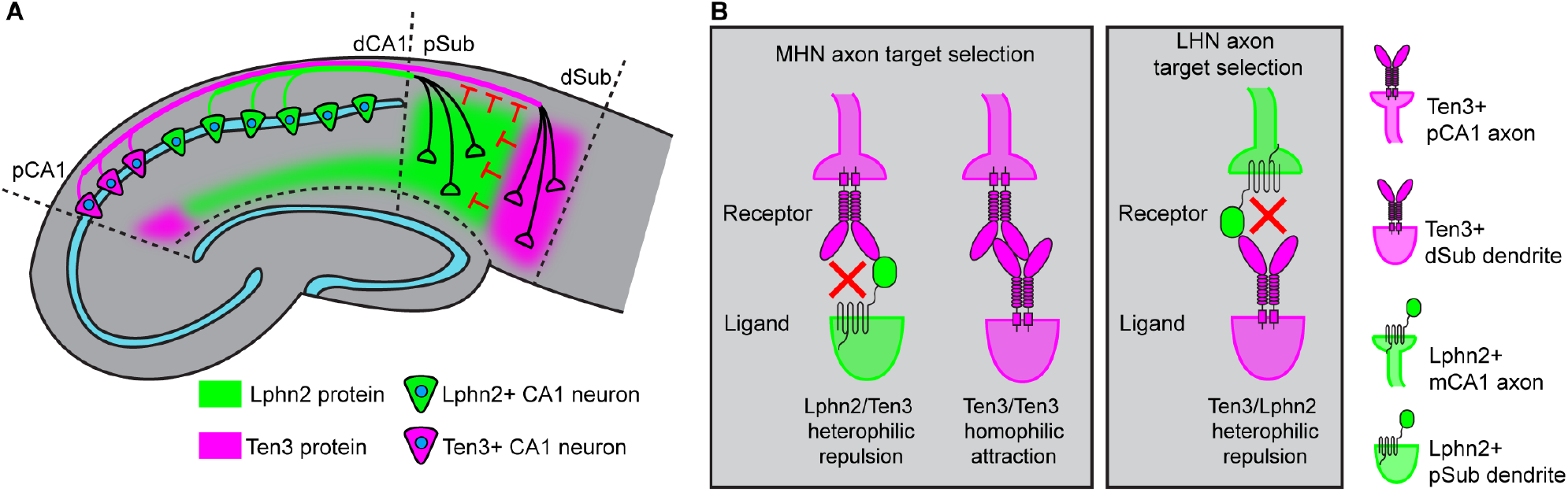
Lphn2 and Ten3 instruct target selection of hippocampal axons through reciprocal repulsions. (**A**) Ten3+ pCA1 axons target Ten3+ dSub via repulsion from Lphn2 and attraction to Ten3 in the subiculum. Lphn2+ mCA1 axons target Lphn2+ mSub via repulsion from Ten3 in dSub. (**B**) High-magnification view of ligand–receptor interactions that instruct target selection of Ten3+ MHN (left) and Lphn2+ LHN (right) axons. Red crosses symbolize repulsion.

Our results have expanded our understanding of the roles of Lphn2 by showing that it acts both cell non-autonomously in targets and cell autonomously in axons during the target selection stage of hippocampal circuit assembly, preceding synapse formation. This is in contrast to previous studies suggesting that Latrophilins act strictly as postsynaptic adhesion molecules to establish or maintain synapses (Anderson et al., 2017; Sando et al., 2019; Südhof, 2018). While defects in axon targeting may contribute to previously reported findings of synaptic deficits in *Latrophilin* early postnatal loss-of-function experiments (Anderson et al., 2017; Sando et al., 2019), our study does not rule out the possibility that Latrophilin/Teneurin interactions play additional roles in synaptic adhesion if the repulsive mechanism is switched off after target selection is complete. Latrophilins bind both Teneurins and FLRTs, and the cooperative binding of these three proteins has been implicated in directing synapse specificity and repulsion-mediated neuron migration (Del Toro et al., 2020; Sando et al., 2019). However, misexpression of a mutant Lphn2 that cannot bind FLRT (Figure S5E and F) still repelled Ten3+ pCA1 axons to the same extent (Figure 2K), suggesting that FLRT binding is not required for Lphn2/Ten3-mediated repulsion during target selection of axons.

We identify Ten3/Lphn2 as a membrane-bound repulsive ligand/receptor pair during circuit assembly, joining the likes of Ephrin/Eph, Dscam/Dscam, and clustered protocadherin/protocadherin (Cheng et al., 1995; Drescher et al., 1995; Lefebvre et al., 2012; Matthews et al., 2007). The mechanisms of how an extracellular adhesive interaction leads to repulsion are poorly understood. *In vitro* studies have suggested that proteolytic cleavage (Drescher et al., 1995) or endocytosis (Hattori et al., 2000; Zimmer et al., 2003) of the receptor complex may be involved in allowing repulsion after Ephrin/Eph extracellular interactions. Both Latrophilins and Teneurins have been shown to undergo cleavage of their extracellular domains (Araç et al., 2012; Vysokov et al., 2018). It is conceivable that Lphn2-Ten3 binding, while sending a repulsive signal to Ten3+ or Lphn2+ growth cones, also triggers the cleavage of Ten3 and/or Lphn2, thus terminating a transient axon-target interaction.

Our results highlight the cooperation of repulsion and attraction and multifunctionality of cell-surface proteins during target selection. Cooperation of attraction and repulsion has been described at different stages of neuronal circuit assembly (Kolodkin and Tessier-Lavigne, 2011; Sanes and Yamagata, 2009). For example, the PlexB receptor interacts with Sema2a and Sema2b through repulsion and attraction, respectively, to mediate distinct guidance functions during *Drosophila* sensory circuit assembly (Wu et al., 2011). We found that target selection of pCA1 axons is determined by Lphn2/Ten3-mediated repulsion from pSub and Ten3/Ten3-mediated attraction to dSub (Figure 3). Thus, Ten3 acts as a receptor for both repulsive and attractive ligands in the same axon during target selection. Conversely, Ten3 acts as a ligand as an attractant for Ten3+ axons, but a repellent for Lphn2+ axons (Figure 5).

A striking finding in this study is the complementary expression of Ten3 and Lphn2 across all interconnected regions of the hippocampal network (Figure 1I). This is reminiscent of Ephrin-A/EphA counter-gradients found across interconnected regions of the developing visual system (Lambot et al., 2005) that utilize bi-directional Ephrin-A/EphA interactions for the formation of topographic projections (Cang et al., 2008; Egea and Klein, 2007; Feldheim et al., 2000, 2004; Frisén et al., 1998; Rashid et al., 2005). The patterns of Ten3 and Lphn2 expression across the hippocampal network follow a ‘Ten3→Ten3, Lphn2→Lphn2’ rule (Figure 1I). We have demonstrated that target-derived Lphn2 acts as a repulsive ligand for Ten3+ axons in the MHN, and that target-derived Ten3 acts as a repulsive ligand for Lphn2+ axons in the LHN. These reciprocal repulsions may guide target selection across seven additional projections of the MHN and LHN that follow the ‘Ten3→Ten3, Lphn2→Lphn2’ rule. It will be interesting to determine if complementary expression of cell-surface molecules and reciprocal repulsions are used in additional parallel networks across the brain.

The repeated use of the same molecules to guide target selection across extended networks, together with the multifunctionality of a single protein serving as receptor and ligand, contribute to explaining how a limited number of cell-surface molecules can specify a much larger number of connections in the mammalian brain.

## Acknowledgements

We thank T. Südhof for the *Lphn2^mVenus^* and *Lphn2^fl^* mice, the Neuroscience Gene Vector and Virus Core at Stanford University for producing viruses, D. Berns for advice, inspiration, and artwork (Figure 1a, i), J. Ferguson for artwork (Figure 2a and Figure 4k), J. Kebschull for custom MATLAB code, H. Meng for virus preparation, members of the Luo lab for advice and support, D. Berns, J. Kebschull, A. Khalaj, H. Li, J. Li, T. Li, C. McLaughlin, K. Shen and A. Shuster for critiques of the manuscript.

## Author contributions

D.T.P. performed all the experiments and analyzed the data, except for single-cell sequencing sample collection and data processing which was performed by J.H.L., with support from S.R.Q. E.C.G. and C.X. assisted in tissue processing. M. J. W. generated MATLAB code and analyzed data. Y.L. and Z.H. produced custom lentivirus. L.L. supervised the study. D.T.P., J.H.L., and L.L. wrote the paper.

## Supplementary Materials

Materials and Methods

Figures S1–S10

Tables S1

Additional References

