## Supplementary Materials for "Reciprocal repulsions instruct the precise assembly of parallel hippocampal networks"

**This PDF file includes:**

Materials and Methods
Supplementary Text
Figures. S1 to S10
Tables S1

### Materials and Methods

Statistical analysis

Statistical tests were performed using GraphPad Prism 7. No statistical methods were used to determine sample size. Due to the low probability of correctly matching P0 lentivirus and adult AAV injections, all breeder pairs were homozygous, and therefore the experiments were not randomized. Images of aggregation assay (Figure S5), double AAV tracing (Figure S4) and Ten3 and Lphn2 immunostaining are representative of at least three individual experiments. For *Lphn2* gain-of-function experiments (Figure 2), results from all experiments are shown in Figure S6 and S7. See 'Image and data analysis for CA1 axon tracing' section for inclusion criteria.

Mice

All procedures followed animal care and biosafety guidelines approved by Stanford University's Administrative Panel on Laboratory Animal Care and Administrative Panel on Biosafety in accordance with NIH guidelines. Both male and female mice were used, and mice were group housed with access to food and water *ad libitum*. Sequencing experiments used mice heterozygous for *Vglut1-Cre* (Harris et al., 2014) (C57BL/6, and 129 background) and *Ai14* (Madisen et al., 2010) (*Rosa-CAG-LSL-tdTomato-WPRE*, C57BL/6J background) for sorting tdTomato+ excitatory neurons. CD-1 mice from Charles River Laboratories were used for double *in situ* hybridization, Lphn2 overexpression, and double AAV tracing studies. $Ten3^{-/-}$ mice (Leamey et al., 2007) were maintained on the CD-1 background. $Lphn2^{mVenus}$ and $Lphn2^{fl/fl}$ mice (Anderson et al., 2017) were on a C57BL/6 and 129 mixed background and were backcrossed once onto the CD-1 background to improve pup survival rate. $Ten3^{fl/fl}$ mice (Berns et al., 2018) were on a mixed CD-1, C57BL/6, and 129 background and were backcrossed once onto a CD-1 background to improve pup survival rate. All genotyping was performed as previously described (Anderson et al., 2017; Berns et al., 2018; Leamey et al., 2007). The total number of mice injected and screened for each experiment is as follows: Figure 2: *LV-GFP*, 131; *LV-GFP-Lphn2*, 84; *LV-GFP-Lphn2_ΔLec*, 126, *LV-GFP-Lphn2_4A, 71* ; Figure 3: $Lphn2^{+/+}$, 176, $Lphn2^{fl/fl}$, 126, $Lphn2^{fl/fl};Ten3^{fl/fl}$, 112. Figure 4: $Ten3^{+/+}$, 87, $Ten3^{fl/fl}$, 74, $Lphn2^{+/+}$, 71, $Lphn2^{fl/fl}$, 95; and Figure S9: $Lphn2^{+/+};Ten3^{-/-}$, 30, $Lphn2^{fl/fl};Ten3^{-/-}$, 40.

Single-cell RNA-sequencing

To express tdTomato in excitatory projection neurons for sequencing, mice heterozygous for *Vglut1-Cre* and *Ai14* were sacrificed at P8 for single cell isolation and sequencing. The following numbers of mice were used for each dissected regions: pCA1, 6 (3 rounds of isolation, 2 mice each isolation), dCA1, 4 (2 rounds of isolation, 2 mice each isolation), MEC, 2 (1 round of isolation, 2 mice each isolation), LEC, 2 (1 round of isolation, 2 mice each isolation), dSub, 2 (1 round of isolation, 2 mice each isolation) and pSub, 2 (1 round of isolation, 2 mice each isolation). A total of 18 mice including both sexes were used in sequencing experiments.

To isolate tdTomato+ cells for single-cell sequencing, mice were briefly anesthetized with isoflurane and decapitated, and the brain was isolated in ice-cold ACSF (2.5 mM KCl, 7 mM $MgCl_2$, 0.5 mM $CaCl_2$, 1.3 mM $NaH_2PO_4$, 110 mM choline chloride, 25 mM $NaHCO_3$, 1.3 mM sodium ascorbate, 20 mM glucose, 0.6 mM sodium pyruvate, bubbled in 95% $O_2$ / 5% $CO_2$). Brains were embedded in 3% low-melting point agarose (Fisher BP165-25) in ACSF at 37°C, cooled to 4°C, and then cut on a vibratome into 350-µm floating sections. For CA1 and subiculum dissections, sections were cut on the sagittal plane; for entorhinal cortex dissections, sections were cut on the horizontal plane. Next, we visualized the fluorescent tdTomato labeling and used the atlas as a guide, to cut out the regions of interest as accurately as possible. Microdissected tissue was incubated at 37°C in papain enzyme mix + 800 nM kyneurenic acid (Worthington) for 30 minutes and triturated gently with a P200 pipette every 15 minutes thereafter until fully dissociated, usually within 1 hour of total incubation time. The cell suspension was spun down at 350 *g* for 10 min at room temperature, neutralized with ovomucoid inhibitor, spun again, washed in ACSF, stained with Hoechst for 10 minutes (1:2000, Life Technologies: H3570), washed, filtered (Falcon 532235), and resuspended in 2 mL ACSF. FACS was performed using the Sony SH800 system with a 130-µm nozzle suitable for the large size of excitatory neurons. Singlet cells were selected based on low FSC-W, and gated on Hoechst (nuclear stain that penetrates cell membrane) and tdTomato double positivity to identify labeled healthy neurons. Cells fulfilling these criteria were over 100× brighter than background, and were unambiguously identifiable. Single cells were sorted at a low flow rate (<100 events/second), and at the highest purity setting (Single Cell) into 384-well hard shell PCR plates (BioRad HSP3901) containing 0.4 µl lysis buffer [0.5 U Recombinant RNase Inhibitor (Takara Bio, 2313B), 0.0625% TritonX-100 (Sigma, 93443-100ML), 3.125 mM dNTP mix (Thermo Fisher, R0193), 3.125 µM Oligo-dT$_{30}$VN (Integrated DNA Technologies, 5′AAGCAGTGGTATCAACGCAGAGTACT$_{30}$VN-3′) and 1:600,000 ERCC RNA spike-in mix (Thermo Fisher, 4456740)] in each well, respectively. Following FACS, plates were spun down, sealed and stored at –80 °C. cDNA synthesis and library preparation protocols were adapted from the SMART-Seq2 protocol (Picelli et al., 2014) (384-well processing utilized 0.4 µl starting volumes, and will hereafter be referred to as 1 unit. Plates were first thawed on ice followed by primer annealing (72°C, for 3 minutes, then on ice). For reverse transcription, 1.5 units of reaction mix [16.7 U/µL SMARTScribe Reverse Transcriptase (Takara Bio, 639538), 1.67 U/µL Recombinant RNase Inhibitor (Takara Bio, 2313B), 1.67× First-Strand Buffer (Takara Bio, 639538), 1.67 µM TSO (Exiqon, 5′-AAGCAGTGGTATCAACGCAGAGTGAATrGrGrG-3′), 8.33 mM dithiothreitol (Bioworld, 40420001-1), 1.67 M Betaine (Sigma, B0300-5VL) and 10 mM $MgCl_2$ (Sigma, M1028-10X1ML)], was added with a Formulatrix Mantis liquid handler. The reaction was then carried out by incubating wells on a thermocycler (Bio-Rad) at 42°C for 90 min, and stopped by heating at 70°C for 5 min. Subsequently, 3.75 units of PCR mix (1.67× KAPA HiFi HotStart ReadyMix (Kapa Biosystems, KK2602), 0.17 µM IS PCR primer (IDT, 5′-AAGCAGTGGTAT CAACGCAGAGT-3′), and 0.038 U/µL Lambda Exonuclease (NEB, M0262L) was added to each well. PCR was then performed using the following program: 1) 37°C for 30 min, 2) 95°C for 3 min, 3) 21 cycles of 98°C for 20 s, 67°C for 15 s and 72°C for 4 min, and 4) 72°C for 5 min. cDNA from every well was quantified using Quant-iT™ PicoGreen™ dsDNA Assay Kit (Thermo Fisher: P11496), and diluted to 0.4 ng/µL in Tris-EDTA before tagmentation. Before tagmentation, we reformatted the samples into a standardized 384-well format, and used the Formulatrix Mantis and Mosquito (TTP Labtech) to automatically perform all liquid handling steps. Tagmentation was performed on double-stranded cDNA using the Nextera XT Library Sample Preparation kit (Illumina, FC-131-1096). Each well was mixed with 0.8 µL Nextera tagmentation DNA buffer and 0.4 µL Tn5 enzyme, then incubated at 55°C for 10 min. The reaction was stopped by adding 0.4 µL Neutralization Buffer and centrifuging at room temperature at 3,220 *g* for 5 min. Indexing PCR reactions were performed by adding 0.4 µL of 5 µM i5 indexing primer, 0.4 µl of 5 µM i7 indexing primer, and 1.2 µL of Nextera NPM mix. PCR amplification was carried out using the following program: 1) 72°C for 3 min, 2) 95°C for 30 s, 3) 12 cycles of 95°C for 10 s, 55°C for 30 s and 72°C for 1 min, and 4) 72°C for 5 min. After library preparation, wells of each 384-library plate were pooled using a Mosquito liquid handler, and consolidated into one tube. Pooling was followed by two final purifications using 0.8× AMPure beads (Fisher, A63881). Library quality was assessed using capillary electrophoresis on a Fragment Analyzer (AATI), and libraries were quantified by qPCR (Kapa Biosystems, KK4923) on a CFX96 Touch Real-Time PCR Detection System (Bio-Rad). Libraries were sequenced on the NovaSeq 6000 Sequencing Systems (Illumina) using 2 × 100-bp paired-end reads, respectively. Sequences were de-multiplexed using bcl2fastq. Reads were aligned to the mouse mm10 genome (with Cre and tdTomato genes added) using STAR version 2.5.4 (Dobin et al., 2013). Gene counts were produced using HTseq version 0.10.0 (Anders et al., 2015), for only exons, with the 'intersection-strict' flag. The raw data resulting from this was a matrix 4045 cells × 21171 genes.

Single-cell RNA-sequencing data analysis

Before generating the transcriptomic map featured in Figure S1B, we first coarsely analyzed the cell datasets from each dissected region individually (pCA1, dCA1, pSub, dSub, MEC, LEC) to identify cells that should be: 1) removed from the dataset based on the expression of known markers from unrelated contaminating regions such as CA2/3), or 2) unambiguously relabeled as a different region based on marker expression. To do this, we first removed cells if they expressed fewer than 2000 genes, and removed genes if they were detected in fewer than 3 cells. This resulted in a dataset of 3691 cells × 18865 genes. We aimed for our dataset to contain > 500 cells for each subregion, and collected cells in batches until this number was reached. The breakdown of the 3691 cell dataset was pCA1: 701 cells, 3 batches; dCA1: 780 cells, 2 batches; pSub: 500 cells, 1 batch; dSub: 678 cells, 1 batch; MEC: 507 cells, 1 batch; LEC: 525 cells, 1 batch. We next subsetted this data based on the 6 dissection regions, and used default settings for filtering, variable gene selection, dimensionality reduction, and clustering (resolution = 0.5) in Seurat v3.0 (Butler et al., 2018). This resulted in a Seurat object for each dissected region, where cells were visualized using a 2-dimensional t-distributed stochastic neighbor embedding (tSNE) of the PC-projected data, and differential expression marker analysis could be performed to identify cell clusters that should be removed or re-labeled. This resulted in between 5 and 10 clusters for each subregion dataset (Figure S2). To identify marker genes for the clusters in each subregion, we used the FindAllMarkers function in Seurat v3.0 and applied this to each Seurat object. This function performs differential expression analysis (Wilcoxon rank sum) between a specific cluster, compared with all other cells not in the cluster, and then iterates through all clusters. Within Seurat, "PCT" refers to the fraction of cells within a population expressing a specific gene. We considered only marker genes that were expressed in at least 25% of cells in either of the comparison groups (min.pct = 0.25), and exhibited a log fold change threshold > 0.25, and plotted the top 10 genes for each cluster using the DoHeatmap function (Figure S2) for visualization. To characterize the anatomical identity of each cluster, we checked the expression pattern of the top 10 genes using *in situ* hybridization images from Allen Institute website (www.mouse.brain-map.org) (Lein et al., 2007). Those clusters which were enriched for genes not in CA1, subiculum or entorhinal cortex were removed (i.e., CA2/3 and presubiculum). Those clusters which were enriched for genes not from their dissected region but still from CA1, subiculum, or entorhinal were re-classified (i.e., dCA1 cells re-classified as pSub).

Generation of the transcriptomic map in Figure S1B

Following the removal of contaminating cells and the relabeling of cells based on region, we generated the final Seurat object featured in Figure S1B under the following conditions. As before, cells expressing fewer than 2000 genes and genes detected in fewer than 3 cells were already removed. We also removed cells expressing *Gad2* > 0, as these were putative inhibitory neurons. This resulted in a final, high-quality dataset of 3382 cells × 18724 genes, where the median cell expressed 6254 genes, from 759819 reads. Cell counts based on the re-labeled regions were: pCA1: 582, dCA1: 682, pSub: 582, dSub 509, MEC: 505, LEC: 522. To generate the tSNE map in Figure S1B, a Seurat object was generated similar to before with default settings. Of note, we performed the ScaleData function and regressed out the effects of the # of genes and the # of reads, and 11 PCs were used in the FindNeighbors function based on a steep dropoff in variance explained by visual inspection of the elbow plot. Cells were visualized using the DimPlot, FeaturePlot, and VlnPlot functions.

Identification of genes with inverse expression to Ten3

To identify genes with inverse expression to Ten3 across CA1, subiculum, and entorhinal cortex, we first subsetted the total data into three groups based on the updated region labels. Within each region, we next identified cells with expression of Ten3 (*Odz3)* in the top 95$^{th}$ percentile in each region, and labeled these as *Ten3*-HIGH. Conversely, we identified the cells with no expression of Ten3 in each region, and labeled these as *Ten3*-NONE. Thus, for each region (CA1, Sub, EC) we had a set of *Ten3*-HIGH and *Ten3*-NONE cells to compare gene expression. To perform differential expression, we compared *Ten3*-HIGH and *Ten3*-NONE cells in each region using the FindMarkers function in Seurat v3.0, which uses the Wilcoxon rank sum test. Compared to our coarse survey of region-based marker genes, we more stringently considered only genes that were expressed in at least 50% of cells in either of the comparison groups (min.pct = 0.5). We also required that the difference in membership between the two groups ($PCT_{Ten3\text{-HIGH}} - PCT_{Ten3\text{-NONE}}$) for considered genes to be >0.1 (min.diff.pct = 0.1). The genes that fit these criteria are listed in **Supplementary Table 1**.

Double *in situ* hybridization

Mice were injected with 2.5% Avertin and were transcardially perfused with PBS followed by 4% paraformaldehyde (PFA). Brains were dissected and post-fixed in 4% PFA for 1 hour, cryoprotected for about 24 hours in 30% sucrose. Brains were embedded in Optimum Cutting Temperature (OCT, Tissue-Tek), frozen in dry ice cooled isopentane bath and stored at –80°C until sectioned. *In situ* hybridizations were performed as previously described (Weissbourd et al., 2014) with the following modifications. 16-µm sections were collected on Superfrost Plus slides and used for fluorescent *in situ* hybridization. *Ten3* and *Lphn2* probes were amplified using primer pairs F-5'-GTGGCTAAAAGCCCACTGTTGCC-3', R-5'-GAATGGCCCACTGACCTCGCG-3' and F-5'-ACCAGTAGCAATCAAGCCACA-3', R-5'-AAGGACCACCTTGGTCGAGT-3', respectively. PCR products were cloned into *pCR4-TOPO* (K457502) and RNA probes were transcribed using T3 or T7 RNA polymerases. *Ten3* probe was labeled with fluorescein (Roche 11685619910) and *Lphn2* probe was labeled with DIG (Roche 11277073910). Slides were hybridized, washed, blocked and incubated with alkaline phosphatase-conjugated anti-DIG-AP antibody (1:1,000, Roche 11093274910) and anti-Fluorescein-HRP (1:200, Perkin Elmer NEF710001EA). Sections were developed with TSA-Plus Fluoroscein system (NEL741001KT) for 15 minutes, 3 hrs with Fast Red TR/Naphthol As-MX (Sigma F4523), counterstained with DAPI, washed and mounted with Fluoromount-G (SouthernBiotech). Sections were imaged using the tile scan function on Zeiss LSM 780 confocal microscope (10x magnification). Fluorescence intensity measurements on unprocessed images were taken using FIJI and data processing was performed using MATLAB. For CA1 measurements, a 75-pixel-wide segmented line was drawn along the cell body layer from proximal to distal. pCA1 was determined by increased DAPI intensity versus CA2 and dCA1 was determined by the condensed CA1 cell body layer disappearing. For subiculum quantification, a 200-pixel-wide segmented line was drawn through the cell body layer from proximal to distal. pSub was defined by the region adjacent to the CA1 cell body layer and dSub was defined as the regions directly before the cell dense presubiculum. For entorhinal cortex, a 250-pixel-wide segmented line was drawn from medial to lateral. Segmented lines were drawn, straightened using the Straighten function, background subtraction was performed using the Subtract function and intensity values were measured using the Plot Profile command (performed in FIJI). The intensity plots were resampled into 100 equal bins (MATLAB) and each individual trace was normalized to a maximum of 100. Traces from each mouse were averaged, normalized to a maximum of 100 again and then the normalized average trace from each mouse was combined to give the final distribution (Figure 1 and Figure S3). Numbers of sections used for quantifications are as follows: horizontal section CA1: 10, 9, 10 sections from 3 mice; horizontal section subiculum: 10, 10, 11 sections from 3 mice; horizontal section entorhinal cortex: 5, 9, 8 sections from 3 mice; sagittal section CA1: 8, 8, 4 sections from 3 mice; and sagittal section subiculum: 8, 7, 4 sections from 3 mice.

Immunostaining

Mice were injected with 2.5% Avertin and were transcardially perfused with PBS followed by 4% paraformaldehyde (PFA). Brains were dissected and post-fixed in 4% PFA for 1 hour (for P8 Ten3 and Lphn2 staining) or 4–6 hours (for adult GFP and mCh staining), cryoprotected for about 24 hours in 30% sucrose. Brains were embedded in Optimum Cutting Temperature (OCT, Tissue-Tek), frozen in dry ice cooled isopentane bath and stored at –80°C until sectioned. 60-μm thick floating sections were collected in PBS + 0.02% sodium azide and stored at 4°C. Sections were incubated in the following solutions at room temperature unless indicated: 1 hour in 0.3% PBS/Triton X-100 and 10% normal donkey serum, two nights (adult GFP and mCh staining) or four nights (P8 Ten3 and Lphn2 staining) in primary antibody at 4°C in 0.3% PBS/Triton X-100 and 10% normal donkey serum, 3 × 15 min in 0.3% PBS/Triton X-100, overnight in secondary antibody + DAPI (1:10,000 of 5 mg $ml^{-1}$, Sigma-Aldrich) in 0.3% PBS/Triton X-

100 and 10% normal donkey serum, 2 × 15 min in 0.3% PBS/Triton X-100, and 15 min in PBS. Sections were mounted with Fluoromount-G (SouthernBiotech). Primary antibodies used were rabbit anti Ten3 (1:500) (Berns et al., 2018), chicken anti-GFP (1:2,500, Aves Labs, GFP-1020) and rat anti-mCherry (1:1,000, Thermo Fisher, M11217). Secondary antibodies conjugated to Alexa 488, Alexa 568, or Cy3 (Jackson ImmunoResearch) were used at 1:500 from 50% glycerol stocks. For quantification of Lphn2 and Ten3 protein images were taken with a Zeiss LSM 780 confocal microscope (10x magnification) and quantification was done as described for *in situ* hybridization except fluorescence intensity measurements were taken from the molecular layers of CA1 (150 pixels wide) and subiculum (150 pixels wide) and layer III of the entorhinal cortex (250 pixels wide). Numbers of sections used for quantifications are as follows: horizontal CA1: 10, 7, 6 sections from 3 mice; horizontal subiculum: 11, 7, 4 sections from 3 mice; horizontal entorhinal cortex: 5, 4, 5 sections from 3 mice; sagittal CA1: 10, 10 sections from 2 mice; and sagittal subiculum: 10, 10 sections from 2 mice. Sections for pCA1 axon quantification in subiculum were imaged using a Zeiss epifluorescence microscope.

<u>Aggregation assay</u>

K562 cells (ATCC, CCL-243) were grown in RPMI-1640 + Glutamax (Gibco, 61870127) with 10% FBS (Gibco A3160501) and 1× penicillin/streptomycin (Gibco 15140122). K562 cells were electroporated with Neon Transfection System 100-µl tips (Thermo Fisher MPK5000) using 1450 V, 10 ms, and 3 pulses. The following amounts of dual reporter plasmid (gene of interest and fluorescent reporter in the same construct) were used: 15 µg of *Ef1α-Ten3 CMV–GFP*, 10.9 µg of *Ef1α-Lphn2*;*CMV–mCh*, 5 µg *Ef1α-Empty*;*CMV–mCh* and 5 µg *Ef1α- Empty;CMV–GFP*. For each electroporation, 2 ×$10^6$ cells were washed in PBS, resuspended in 120 µl of buffer R, combined with DNA and electroporated. Cells were added to 5 ml of pre-warmed RPMI-1640 with 10% FBS and incubated at 37°C and 5% $CO_2$ for ~20 hours. Cells were collected and resuspended in RPMI-1640 + 10% FBS and incubated for 15 min at 37 °C with 0.1 mg/ml DNase (Worthington LS002060). Cells were resuspended in aggregation media (Neurobasal-A, 4% B27, 2 mM glutamine, 10% FBS, 20 mM HEPES) and passed through a 40-µm cell strainer. 2 ×$10^5$ cells of each condition (4 ×$10^5$ total) were added to wells of a 24-well plate in 1 ml of aggregation media and cells were left to aggregate for 1–2 hours on a nutator at 37°C. Cells were transferred to 2 ml of PBS in a 12-well plate and aggregates were imaged with a Thermofisher Scientific EVOS M500 microscope at 10x magnification. GFP and mCherry channels were combined, thresholded and Analyze Particles (ImageJ) to quantify the aggregate size and number. In each condition the size of a large single cell was calculated and all particles below this value were deleted to filter out the majority of single cells. Particle sizes from three separate experiments were combined and a Kruskal-Wallis test followed by Dunn's multiple comparison was used to test significance between conditions. *Ef1α-Ten3 CMV–GFP*, *Ef1α-Empty CMV–mCh* and *Ef1α- Empty CMV–GFP* were previously generated (Berns et al., 2018) and *Ef1α-Lphn2 CMV–mCh* was generated by using Gibson assembly cloning kit (NEB E5510S) to insert *Lphn2* ORF into *Ef1α- Empty CMV–mCh*. Full length *Lphn2* was amplified from cDNA isolated from P8 mouse hippocampus. The *Lphn2_ΔLec* and *Lphn2_4A* mutations were made using Q5 mutagenesis (NEB, E0552S). The *Lphn2_ΔLec* mutant had amino acids 49-129 removed and the *Lphn2_4A* mutant had amino acids 254, 256, 257 and 313 replaced with alanine. The *Ten3* isoform used contained both NHL and EGF domain alternatively spliced exons (the $A_1B_1$ isoform (Berns et al., 2018)), which is the highest expressed isoform during the development of the CA1→Sub axon projection (Berns et al., 2018).

Lentivirus production

LV-UbC-GFP-Cre was generated by the Neuroscience Gene Vector and Virus core, Stanford University. All other lentivirus constructs were made by inserting either GFP, Lphn2 or Lphn2_ΔLec into the LV-UbC plasmid with a P2A sequence between the two ORFs. GFP was amplified from LV-UbC-GFP-Cre and full length Lphn2 was isolated from P8 mouse hippocampus cDNA. GFP and Lphn2 were inserted into LV-UbC with Gibson assembly cloning kit (NEB E5510S). The Lphn2_ΔLec and Lphn2_4A mutations were made using Q5 mutagenesis (NEB, E0552S). All plasmids were sequenced verified before virus was produced. All custom lentiviruses were generated by transfecting 30 10-cm plates (HEK293T) with 4 plasmids (4.1R, RTR2, VSVg, and transfer vector containing gene of interests). Medium was collected 48 hrs later and centrifuge at 7000 rcf for 18 hrs at 4°C. Viral pellets were dissolved with PBS and further purified with a HiPrep Q FF (16/10) ion-exchange column (GE healthcare) ***(Kinoshita et al., 2012)***. The final concentration of individual lentivirus ranged from $1 \times 10^{11}$ to $6 \times 10^{12}$ infectious copies per ml.

Stereotactic injections in neonatal mice

P0 mice were anesthetized using hypothermia. Subiculum injections were 1.1–1.2 mm lateral, 0.3–0.35 mm anterior and 0.85 mm ventral from lambda and CA1 injections were 1.15 mm lateral, 0.95 mm anterior and 0.8 mm ventral from lambda. 100 nl of lentivirus was injected at 100 nl $min^{-1}$ at the following titers: *LV-UbC-GFP* ($6 \times 10^{12}$ infectious copies per ml), *LV-UbC-GFP-P2A-Lphn2-FLAG* ($6 \times 10^{11}$–$5 \times 10^{12}$ infectious copies per ml), *LV-UbC-GFP-P2A-Lphn2_ΔLec -FLAG* ($1 \times 10^{11}$–$4 \times 10^{11}$ infectious copies per ml), *LV-UbC-GFP-P2A-Lphn2_4A-FLAG* ($1 \times 10^{13}$ infectious copies per ml), *LV-UbC-GFP-Cre* ($1.3 \times 10^{8}$–$5.9 \times 10^{8}$ infectious copies per ml, Neuroscience Gene Vector and Virus core, Stanford University). 200 nl was injected into *Lphn2$^{fl/fl}$;Ten3$^{fl/fl}$* mice.

Stereotactic injections in adult mice

Injections of *AAV8-CaMKIIa-ChR2-mCh* ($2 \times 10^{12}$ infectious copies per ml, Neuroscience Gene Vector and Virus core, Stanford University), *AAV8-EF1a-ChR2-mCh* ($2 \times 10^{12}$ infectious copies per ml, Neuroscience Gene Vector and Virus core, Stanford University) and *AAV8- CaMKIIa-ChR2-YFP* ($2 \times 10^{12}$ infectious copies per ml, Neuroscience Gene Vector and Virus core, Stanford University) were performed at about P36. Mice were anesthetized using isoflurane and mounted in stereotactic apparatus (Kopf). Coordinates for pCA1 were 1.4 mm lateral and 0.9–1.25 mm posterior from bregma, and 1.0–1.12 mm ventral from brain surface; mCA1 were 1.4 mm lateral and 1.55 mm posterior from bregma, and 0.9 mm ventral from brain surface dSub were 3.4 mm lateral, 0 mm anterior, 2.0 mm ventral; msub were 3.6 mm lateral, 0.2 mm anterior, 2.0 mm ventral. Virus was iontophoretically injected with current parameters 5 μA, 7 s on, 7 s off, for 2 min, using pipette tips with an outside perimeter of 10–15 μm. Mice were perfused about 2 weeks later and processed for immunostaining as described above.

Image and data analysis for CA1 axon tracing

Mice were only included if they passed the following criteria: (1) AAV injection site must be in pCA1 (most proximal 30%) or mCA1 ( ~40%–60%) — based on *Ten3* and *Lphn2* mRNA in sagittal sections (Figure S10); (2) lentivirus injections must be in the subiculum, more specifically in dSub for experiments described in Figure 2, in mSub and/or pSub for experiments described in Figure 3A and B, and in both dSub and pSub for experiments described in Figure 3C; (3) pCA1 axons must overlap with lentivirus injection site in subiculum. (4) for mice in Figure 4G and H lentivirus injections sites must be in CA1 and not in subiculum. All mice that fulfilled these criteria are reported in Figure 2 to 4 and figs. S6, S7 and S9, and were included in **quantifications**. Images of injections sites (5× magnification) and projections (10× magnification) were acquired of every other 60-μm sagittal section using a Zeiss epifluorescence scope. Due to variation in injection sites within each mouse, exposure was adjusted for each mouse to avoid saturation. Fluorescence intensity measurements on unprocessed images were taken using FIJI and data processing was performed using MATLAB. For injection site quantification, a 30-pixel-wide segmented line was drawn from pCA1 to dCA1 using DAPI signal as a guide. For projection quantification in subiculum, a 200-pixel-wide segmented line was drawn from pSub to dSub through the cell body layer using only DAPI as a guide. From this point injection site and projection images were processed the same. Segmented lines were straightened using the Straighten function, background subtraction was performed using the Subtract function and intensity values were measured using the Plot Profile command (FIJI). For injections that labeled both CA2 and pCA1, CA2 axons were present near the distal border of CA1 and spilled into pSub. These axons had their intensity set to zero by using area selection and the clear function (FIJI). The intensity plots were resampled into 100 equal bins using a custom MATLAB code. Heatmaps were generated in MATLAB using the 'imagesc' function and three-dimensional mountain plots were generated using the 'surf' function. To quantify average axon intensity in GFP+ and adjacent GFP– regions (Figure 2k), we restricted analysis to the most distal 20% of the subiculum. To determine the GFP+ region we identified the intensity-weighted central row using the summed fluorescence of each row and determined the minimal symmetric window of rows around the central row that encompassed at least 50% of the total intensity in the restricted GFP image. This defined a rectangle in the original image that we designated as the GFP+ region. We then computed the mean fluorescence intensity in this region for the mCh channel. We used the two rows above and below (lateral and medial) the designated GFP+ region as the adjacent GFP– region and computed the mean mCh fluorescence across these four rows. To determine mCh fluorescence differences in GFP+ versus GFP– regions, we divided the mCh intensity in the GFP+ region by the mCh intensity in the GFP–region for each mouse (i.e,. GFP+/GFP–). mCh fluorescence intensity GFP+/GFP– was compared across groups using a one-way ANOVA with Tukey's multiple comparisons test using Prism 7 (GraphPad). For trace quantification in Figure 3, 4 and Figure S9, only sections with GFP-Cre expression in the correct subiculum area were quantified. The three GFP-Cre+ sections with the total most axon labelling were combined by summing the three intensity values at each binned position. The summed intensity trace was then normalized to a peak value of 100. The mean position of the injection sites (Figure S10) was calculated by generating a summed intensity trace as above and then multiplying the intensity value by the bin position, summing across the entire axis, and dividing by the sum of the intensity values. To calculate the fraction of axon intensity (Figure 3, 4 and Figure S9), the “area under curve” was calculated for each trace in each 20% segment using Prism 7 (GraphPad). Fraction of axon intensities were compared using a two-way ANOVA with Sidak’s multiple comparisons test. For trace quantification in Figure 4, where *LV-GFP-Cre* was injected into CA1, the three sections with the highest total axon intensity in subiculum were used for quantification. Representative images (Figure 3, 4 and Figure S6, S7 and S9) were taken using a Zeiss LSM 780 confocal microscope (20× magnification, tile scan, max projection) and all images from each individual mouse were processed identically.

Code availability

Custom MATLAB scripts used in data processing and analysis are available from the corresponding author upon request.

Data availability

All data supporting the findings reported in this study are available from the corresponding author upon reasonable request. The single-cell RNA-seq datasets generated and analyzed in the study will be deposited in NCBI Gene Expression Omnibus.

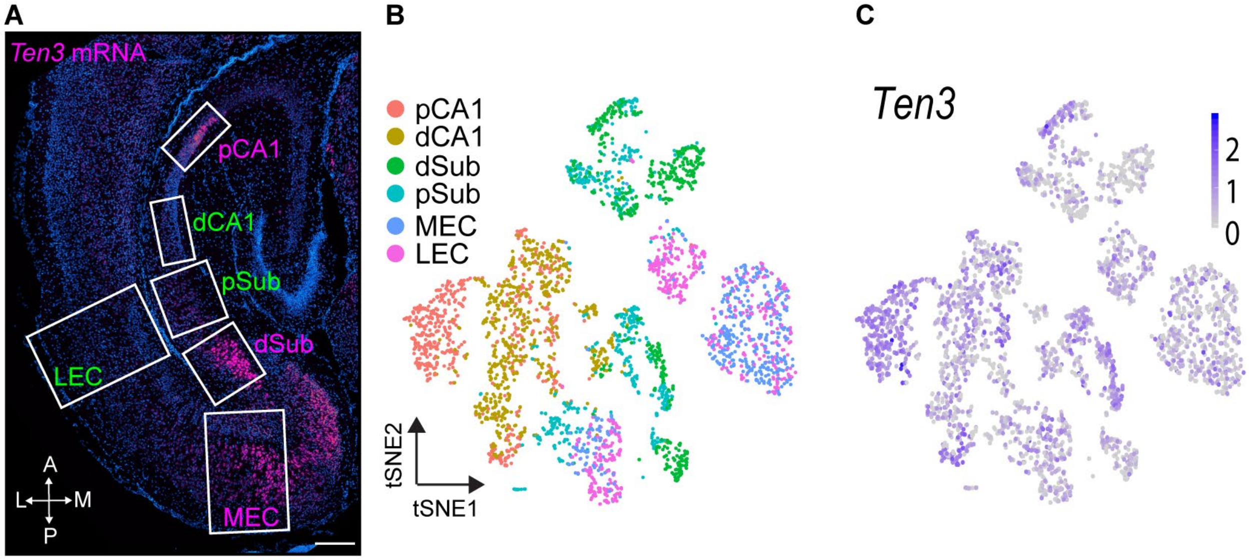


**Figure S1. Identifying genes with expression patterns complementary to *Ten3* using single-cell RNA-sequencing.** (**A**) *In situ* hybridization for *Ten3* on a horizontal P8 mouse brain section with boxes labeling the regions of medial and lateral hippocampal network dissected for sequencing analysis. We micro-dissected boxed regions from *Vglut1-Cre;Ai14* double transgenic mice, dissociated cells, and used fluorescence-activated cell sorting (for tdTomato-positive excitatory neurons) for single-cell RNA sequencing (scRNA-seq) using the Smart-Seq2 platform (Picelli et al., 2014). A, anterior; P, posterior: L, lateral; M, medial. Scale bar, 200 μm. We chose P8 to coincide with the period of target selection for pCA1→dSub axons (Berns et al., 2018). (**B**) tSNE plot of all sequenced cells labelled by dissected region. Identities of regions (color coded) are based on dissection and known molecular markers (Figure S2). Representation of the scRNA-seq data in the tSNE space revealed clusters that segregated partially by subregions but did not clearly delineate the lateral versus medial hippocampal networks. Instead, every cluster contained a combination of cells predominantly from one of the larger regions (CA1, subiculum, and entorhinal cortex). Single cells from contaminating adjacent regions were removed from the dataset or relabeled based on the expression of known marker genes (Figure S2). (**C**) *Ten3* expression represented on tSNE plot in (**B**). We compared the gene expression of cells highly expressing *Ten3* (>95$^{th}$ percentile, *Ten3*-HIGH) with those that did not express *Ten3* (*Ten3*-NONE) within each large region (CA1, subiculum, or entorhinal cortex), tabulating genes with higher average expression in the *Ten3*-NONE group for each region (Table S1).

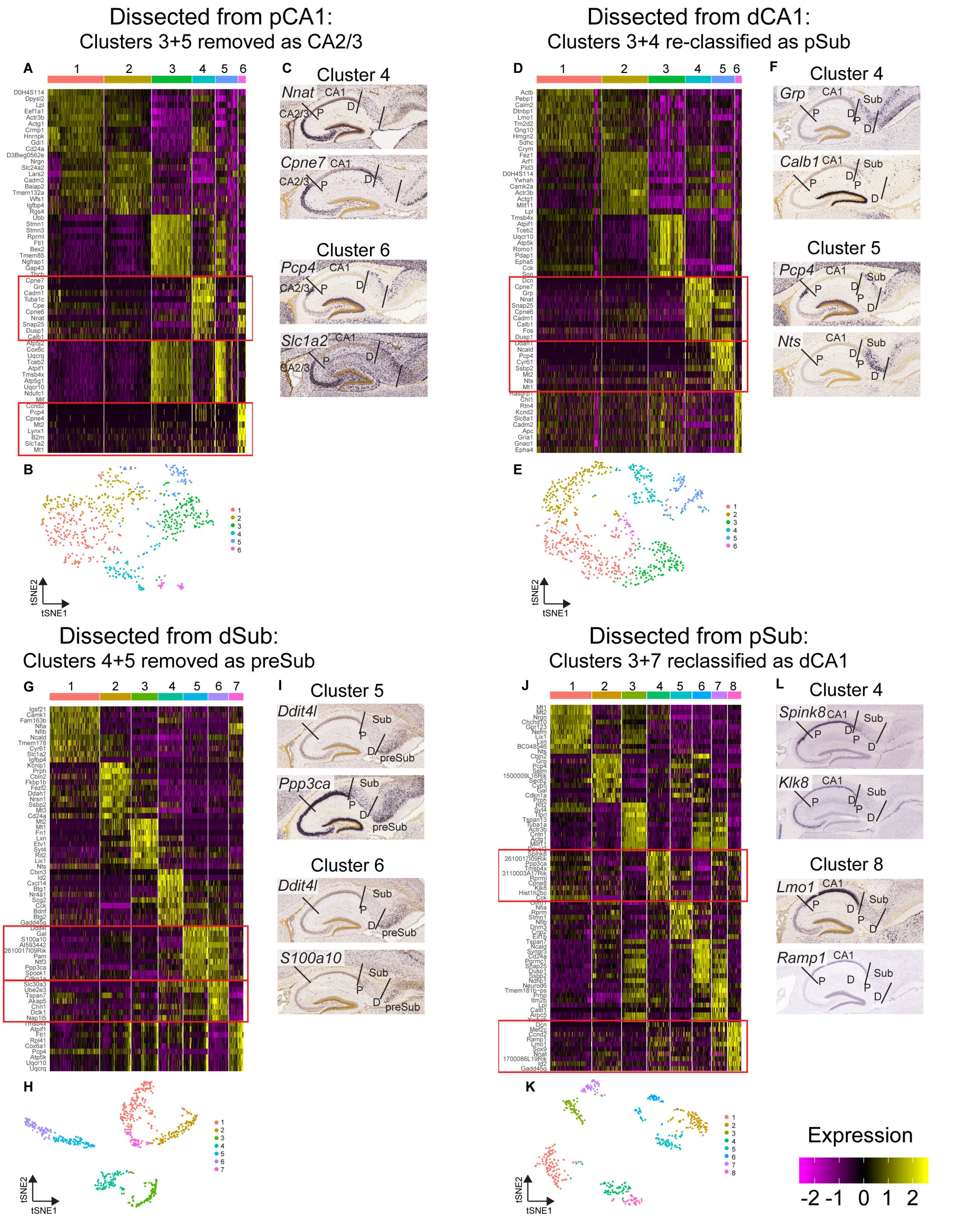


**Figure S2. Removal and re-classification of cells based on molecular markers.** (**A**), (**D**), (**G**), (**J**) Heatmaps highlighting the top 10 differentially expressed genes in each cluster for the dissected regions pCA1, dCA1, dSub, pSub. LEC and MEC were not included here since no cells were removed or re-classified from these regions. Red boxes indicate the clusters that were removed, or re-classified based on molecular markers. (**B**)**,** (**E**), (**H**), (**K**) tSNE plots of all sequenced cells in each dissected region as indicated (see Materials and Methods). (**C**), (**F**), (**I**), (**L**) *In situ* hybridization images from Allen Institute website (www.mouse.brain-map.org) (Lein et al., 2007) highlighting the spatial expression of genes expressed in clusters that were removed or re-classified. We examined the spatial expression pattern of the top 10 genes in each cluster using the *in situ* hybridization images from Allen Institute website (Lein et al., 2007). All clusters except those highlighted (**A**), (**D**), (**G**), and (**J**) expressed molecular markers that corresponded to the region of dissection. In the pCA1 dissection (**A** to **C**), clusters 4 and 6 displayed enriched expression for genes present in CA2/3 and not CA1. Clusters 4 and 6 were thus removed before analysis. In the dCA1 dissection (**D** to **F**), clusters 4 and 5 displayed enriched expression for genes present in subiculum and not CA1, likely due to the close proximity of dCA1 to pSub. Clusters 4 and 5 were thus re-classified as pSub cells before analysis. In the dSub dissection (**G** to **I**), clusters 5 and 6 displayed enriched expression for genes present in presubiculum and not subiculum. Clusters 5 and 6 were thus removed before analysis. In the pSub dissection (**J** to **L**), clusters 4 and 8 displayed enriched expression for genes present in CA1 and not subiculum, likely due to the close proximity of pSub to dCA1. Clusters 4 and 8 were thus re-classified as dCA1 cells before analysis. All *in situ* hybridization images presented here are from P14 mice, except for *Spink8*, *Klk8* and *Ramp1*, which are from P56 mice because there is no data available for these genes at P14.

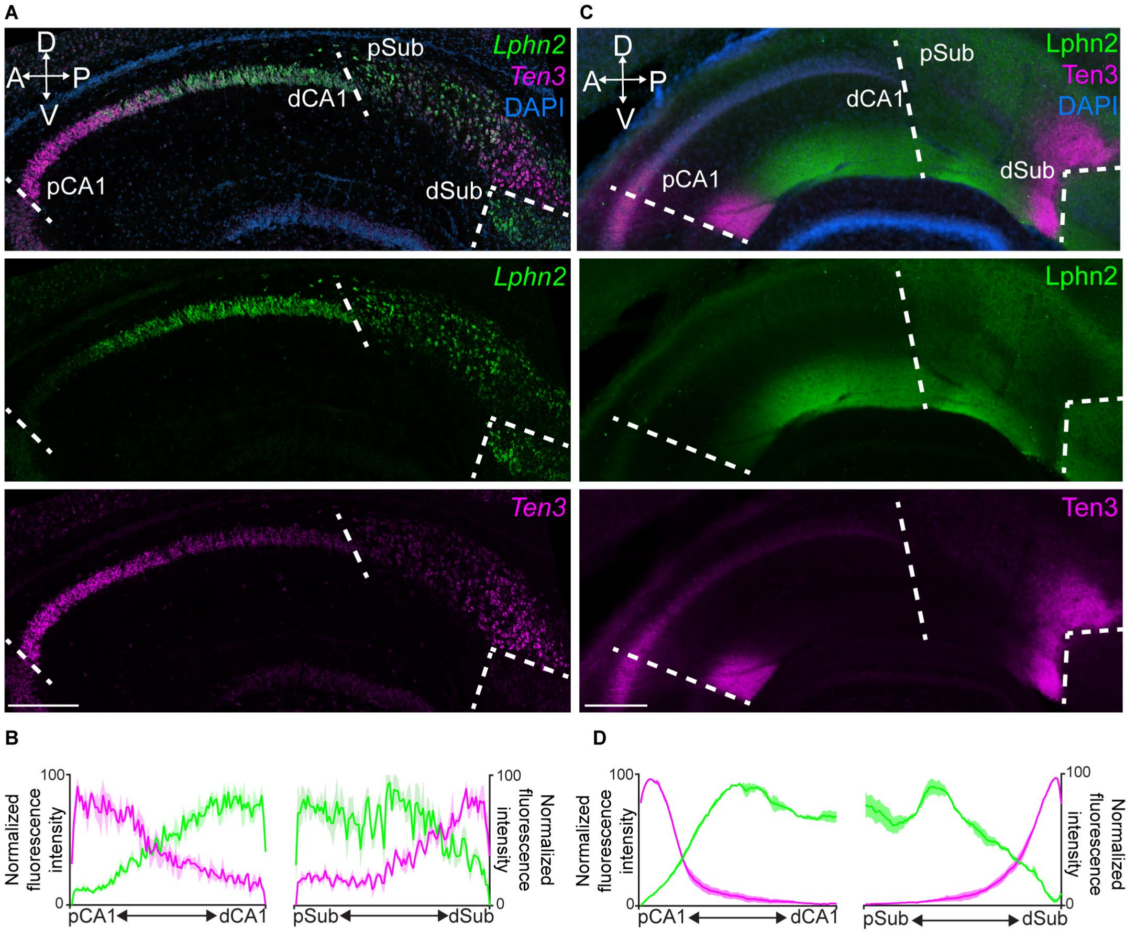


**Figure S3. Complementary expression patterns of *Lphn2* and *Ten3* in sagittal sections.** (**A**) Double *in situ* hybridization for *Lphn2* (middle) and *Ten3* (bottom) mRNA on a sagittal section of P8 mouse brain. Dashed lines represent boundaries between CA1 and subiculum as labeled in the overlay (top). A, anterior; P, posterior: D, dorsal; V, ventral. (**B**) Quantification of *Lphn2* and *Ten3* mRNA across the proximal-distal axis of CA1 ($n$ = 3 mice, 23 sections total) and subiculum ($n$ = 3 mice, 19 sections total) cell body layers. Mean ± SEM. (**C**) Double immunostaining for Lphn2 (middle; anti-GFP antibody) and Ten3 (bottom) on a sagittal section of P8 *Lphn2-mVenus* knock-in mouse **(Anderson et al., 2017)** brain. Dashed lines represent boundaries between CA1 and subiculum as labeled in the overlay (top). (**D**) Quantification of Lphn2 and Ten3 protein across the proximal–distal axis of CA1 ($n$ = 2 mice, 20 sections total) and subiculum ($n$ = 2 mice, 20 sections total). Mean ± SEM. Scale bars, 200 μm.

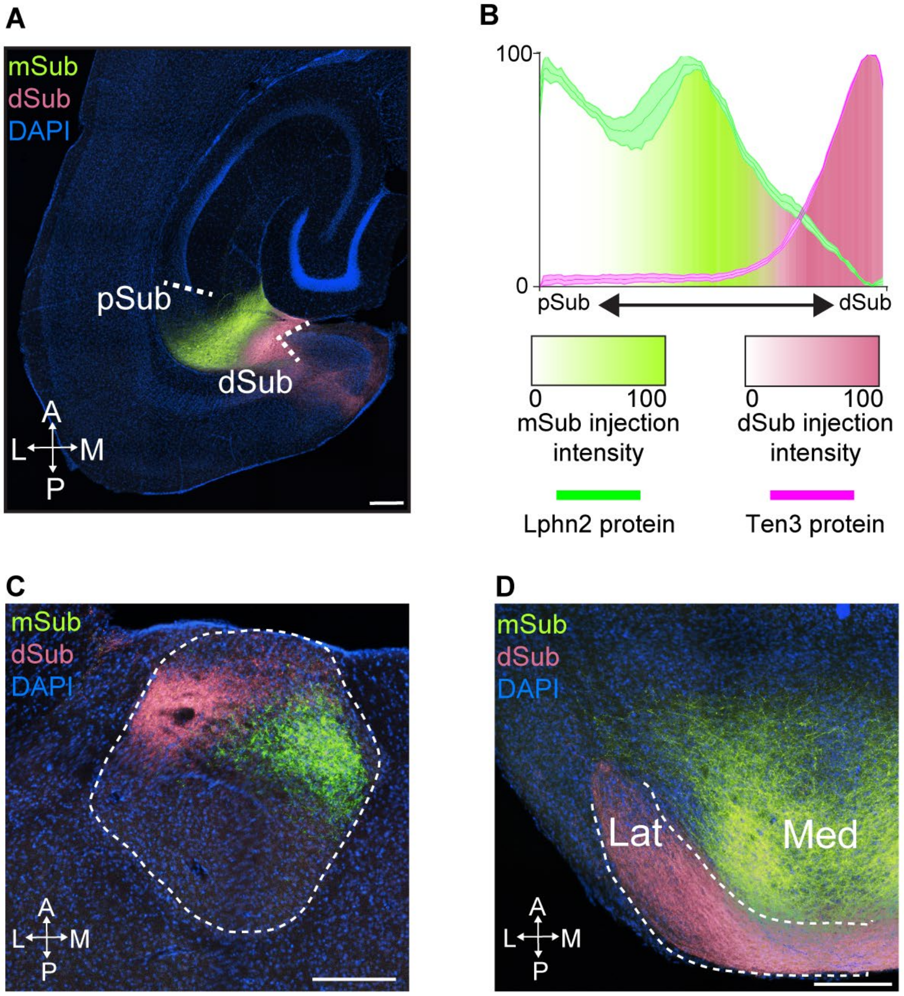


**Figure S4. Anterograde tracing from subiculum.** (A**)** Injection sites of dual anterograde tracing from Lphn2+ mid-subiculum (mSub) and Ten3+ dSub in a P55 mouse brain. (**B**) Quantification of injection sites position along the proximal–distal axis (intensity shown as color gradients) superimposed on Lphn2 and Ten3 protein distribution (curves) as shown in Figure 1G. (**C**) Projection of subiculum axons from (**A**) in the anteroventral nucleus of the thalamus. Lphn2+ axons target the medial region of the anteroventral thalamus, whereas Ten3+ axons target the lateral region. (**D**) Projection of subiculum axons from (**A**) in the medial mammillary nucleus. Lphn2+ axons target the medial regions, whereas Ten3+ axons target the lateral region. A, anterior; P, posterior; L, lateral; M, medial. Scale bars, 200 µm.

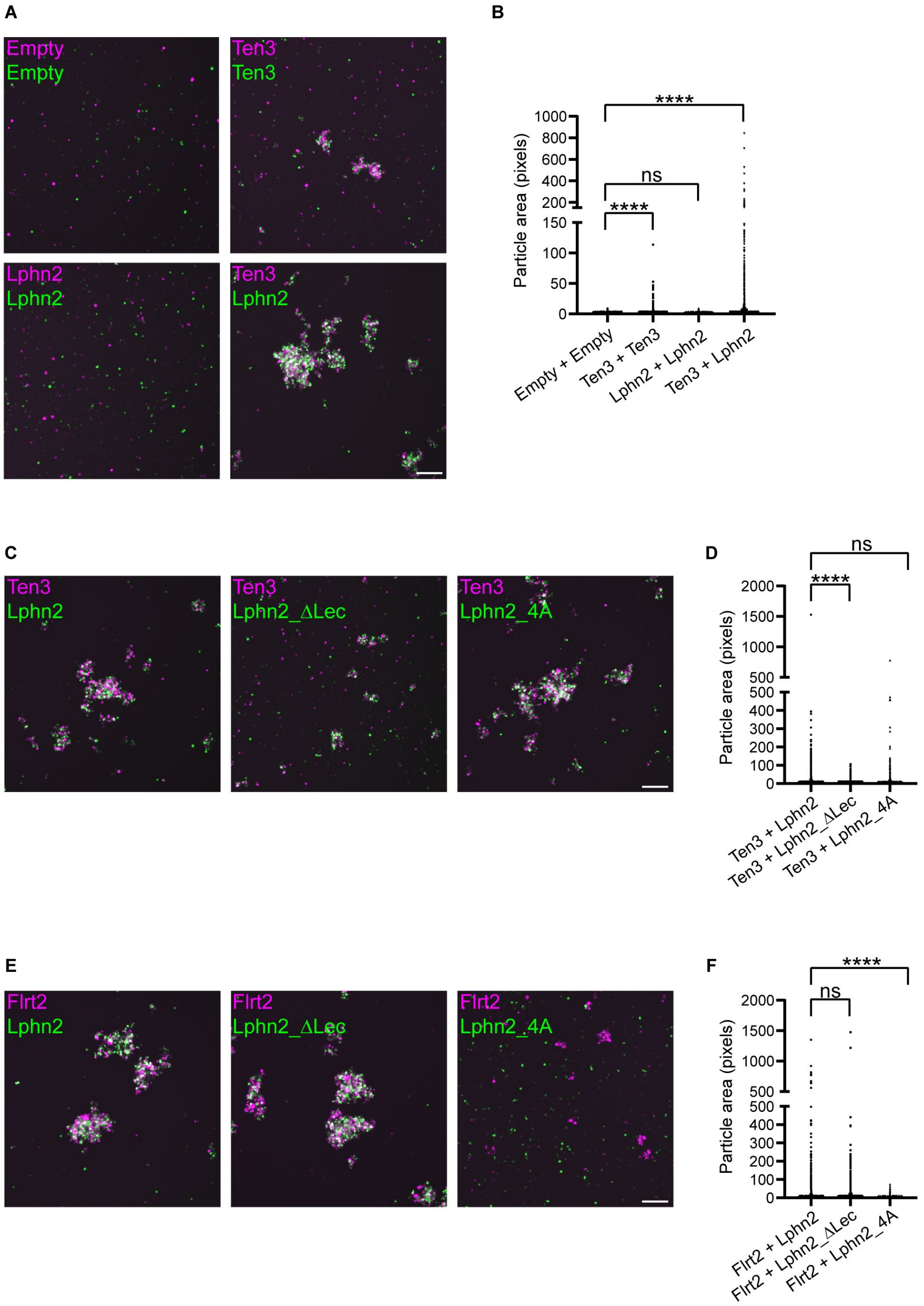


**Figure S5. Aggregation assays to test Ten3 and Lphn2 in promoting *trans*-cellular interactions and to validate mutant Lphn2 that does not bind Ten3 or FLRT2.** (**A**) Top left, non-adherent K562 cells transfected with RFP alone (empty) or GFP alone (empty) do not form aggregates when mixed. Top right, Ten3/RFP-expressing cells and Ten3/GFP-expressing cells form aggregates when mixed. Bottom left, Lphn2/RFP-expressing cells and Lphn2/GFP-expressing cells do not form aggregates when mixed. Bottom right, Ten3/RFP-expressing cells and Lphn2/GFP-expressing cells form aggregates when mixed. (**B**) Quantification of particle area confirms that Ten3 promotes homophilic adhesion (Berns et al., 2018), suggests that Lphn2 does not promote homophilic adhesion, and confirms that Ten3 and Lphn2 promote heterophilic adhesion (Berns et al., 2018; Boucard et al., 2014; Li et al., 2018; del Toro et al., 2020). Mean ± SEM; Kruskal-Wallis test with Dunn's multiple comparison test; ns, not significant; ****P ≤ 0.0001. (**C**) Left, Ten3/RFP-expressing cells and Lphn2/GFP-expressing cells form aggregates when mixed. Middle, Ten3/RFP-expressing cells and Lphn2_ΔLec/GFP-expressing cells form only small aggregates when mixed. Right, Ten3/RFP-expressing cells mixed with Lphn2_4A/GFP-expressing cells form aggregates when mixed. (**D**) Ten3 + Lphn2 aggregates are significantly bigger than Ten3 + Lphn2_ΔLec aggregates but not different from Ten3 + Lphn2_4A aggregates. Thus, Lphn2_ΔLec but not Lphn2_4A disrupts Ten3 binding. Mean ± SEM; Kruskal-Wallis test with Dunn's multiple comparison test; ns, not significant; ****P ≤ 0.0001. (**E**) Left, FLRT2/RFP-expressing cells and Lphn2/GFP-expressing cells form aggregates when mixed. Middle, FLRT2/RFP-expressing cells and Lphn2_ΔLec/GFP-expressing cells form aggregates when mixed. Right, FLRT2/RFP-expressing cells and Lphn2_4A/GFP-expressing cells form small aggregates that only contain FLRT2/RFP-expressing cells. The presence of aggregates that only contain FLRT2/RFP-expressing cells confirms FLRT-mediated homophilic adhesion previously described (Seiradake et al., 2014). (**F**) FLRT2 + Lphn2 aggregates are not significantly different to FLRT2 + Lphn2_ΔLec aggregates but significantly larger FLRT2 + Lphn2_4A aggregates, confirming that Lphn2_4A but not Lphn2_ΔLec disrupts Lphn2-FLRT2 binding. Mean ± SEM; Kruskal-Wallis test with Dunn's multiple comparison test; ns, not significant; ****P ≤ 0.0001. Scale bar, 200 µm.

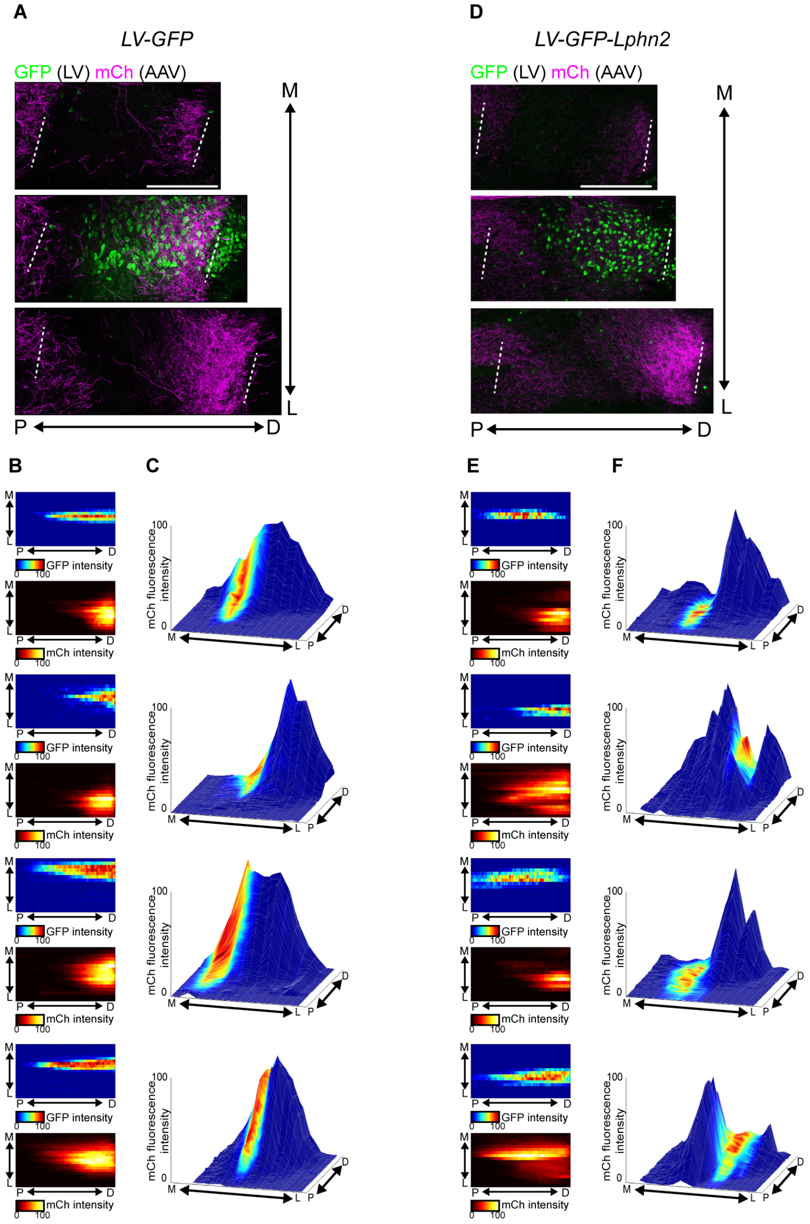


**Figure S6. Additional data related to Lphn2 gain-of-function experiments (Figure 2).** (**A** and **D**) Example images from mice injected with *LV-GFP* (**A**) and *LV-GFP-P2A-Lphn2_FLAG* (**D**) showing pCA1 axons (magenta) and LV (green) in subiculum. Three 60-µm sections that contain pCA1 axons at different positions along the medial–lateral (M–L) axis are shown, with the center one overlapping with the LV injection site. These images correspond to mice analyzed in Figure 2. Scale bars, 200 µm. (**B** and **E**) Heatmaps from additional mice injected with *LV-GFP* (**B**), and *LV-GFP-P2A-Lphn2* (**E**) and analyzed as in Fig 2. (**C** and **F**) Mountain plots from additional mice injected with *LV-GFP* (**C**) and *LV-GFP-P2A-Lphn2* (**F**), and analyzed as in Fig 2.

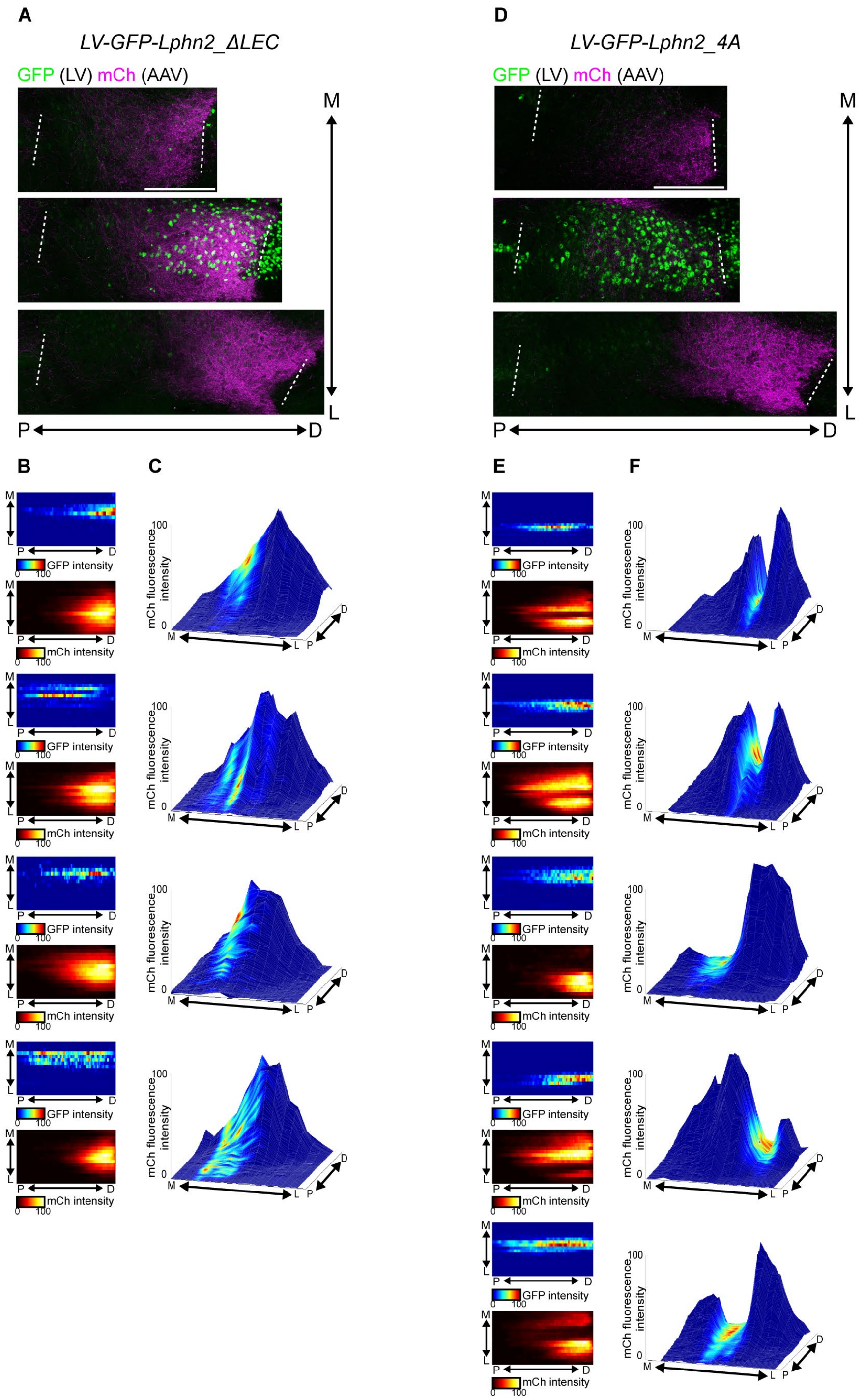


**Figure S7. Additional data related to Lphn2 gain-of-function experiments (Figure 2).** (**A** and **D**) Example images from mice injected with *LV-GFP-P2A-Lphn2_ΔLec_FLAG* (**A**) and *LV-GFP-P2A-Lphn2_4A_FLAG* (**D**), showing pCA1 axons (magenta) and LV (green) in subiculum. Three 60-μm sections that contain pCA1 axons at different positions along the medial–lateral (M–L) axis are shown, with the center one overlapping with the LV injection site. These images correspond to mice analyzed in

Figure 2. Scale bars, 200 μm. (**B** and **E**) Heatmaps from additional mice injected with *LV-GFP-P2A-Lphn2_ΔLec* (**B**) and *LV-GFP-P2A-Lphn2_4A* (**E**) and analyzed as in Figure 2. (**C** and **F**) Mountain plots from additional mice injected with *LV-GFP-P2A-Lphn2_ΔLec* (**C**) and *LV-GFP-P2A-Lphn2_4A* (**F**), and analyzed as in Figure 2.

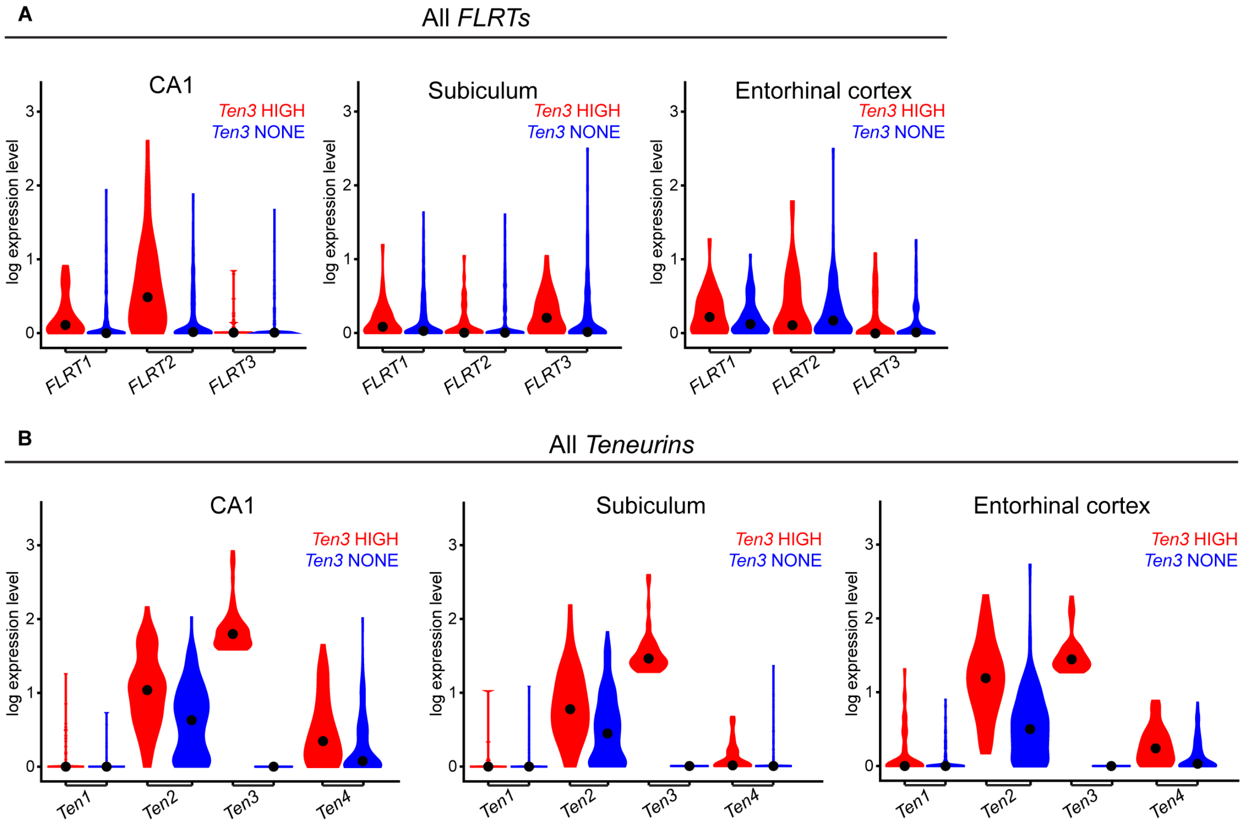


**Figure S8. Expression patterns of *FLRTs* and *Teneurins* in the hippocampal network.** Violin plots of *FLRTs* (**A**) and *Teneurins* (**B**) showing expression in *Ten3*-HIGH and *Ten3*-NONE cells across CA1, subiculum, and entorhinal cortex. Unit of expression level is ln [1+ (reads per 10000)]. Black dots represent the median.

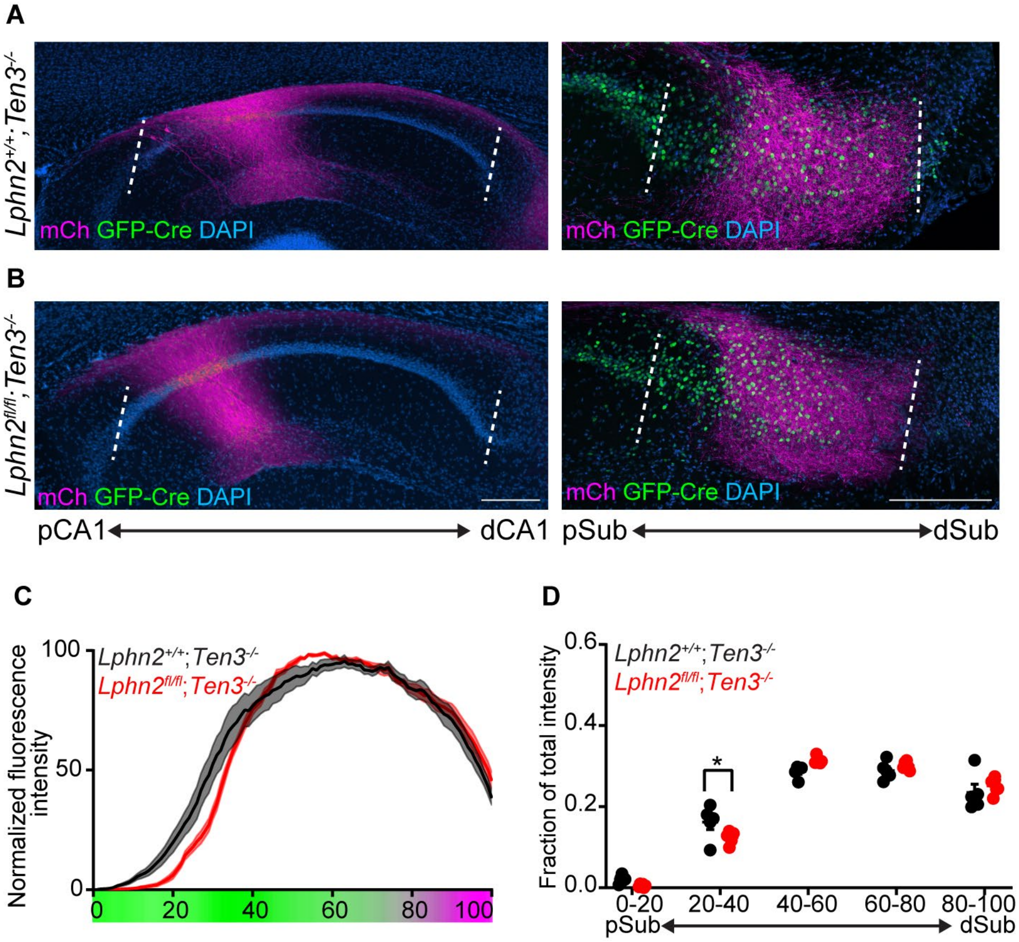


**Figure S9. *Ten3*-null pCA1 axons do not have additional spread into pSub where *Lphn2* is deleted.** (**A** and **B**) Representative images of AAV-mCh (magenta) injections in pCA1 (left) and corresponding projections overlapping with LV-GFP-Cre (green) injection sites in the subiculum (right). Genetic conditions are as indicated. (**C**) Normalized mean fluorescence intensity traces of subiculum projections from pCA1 in GFP-Cre+ sections for *Lphn2*$^{+/+}$*;Ten3*$^{-/-}$ (n = 5 mice) and *Lphn2*$^{fl/fl}$*;Ten3*$^{-/-}$ (n = 6 mice). Mean ± SEM. Color bar under x-axis represents Lphn2 (green) and Ten3 (magenta) expression in subiculum as quantified in Figure S3. (**D**) Fraction of total axon intensity for same data as (**C**) across 20 percent intervals. Mean ± SEM, two-way ANOVA with Sidak's multiple comparisons test, * $P \leq 0.05$. Scale bar, 200 μm. The statistically significant difference at 20–40% is likely because injection sites for *Lphn2*$^{+/+}$*;Ten3*$^{-/-}$ mice were slightly less proximal than for *Lphn2*$^{fl/fl}$*;Ten3*$^{-/-}$ mice (see injection site locations in CA1 in Figure S10).

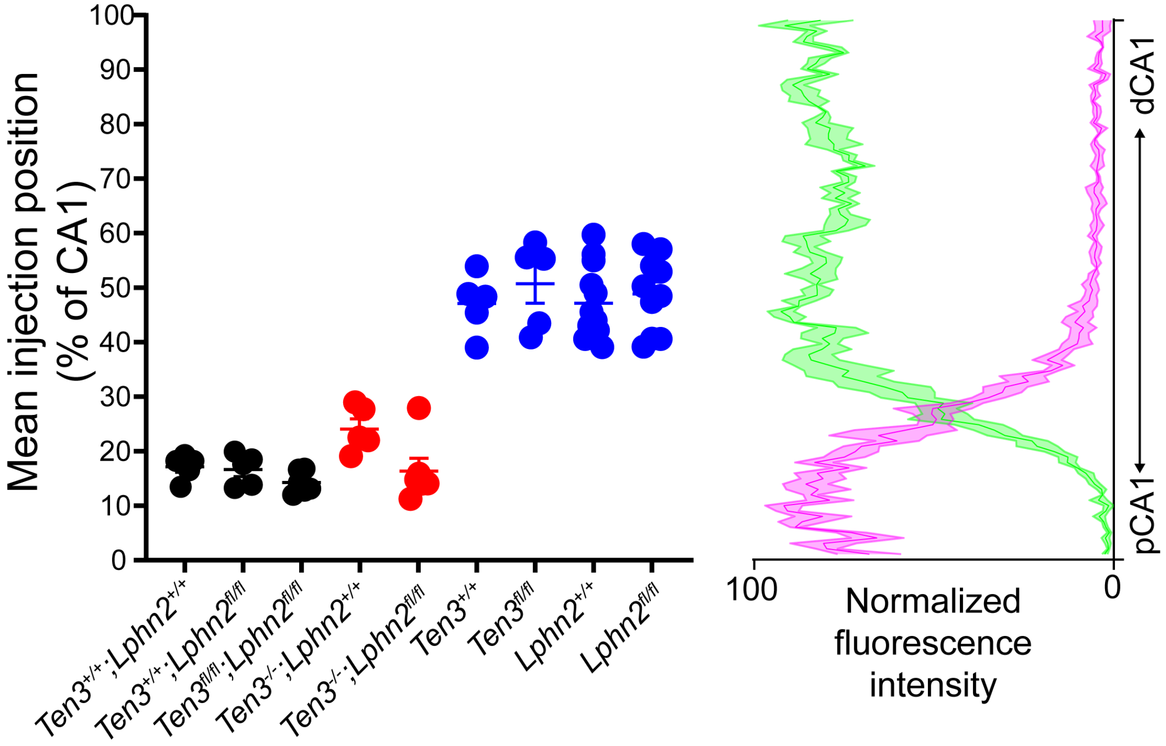


**Figure S10. Mean injection site position of pCA1 injections in mice from Figure. 3, 4, and Figure S9.** All injections sites from mice in Figure 3 (black) and Figure S9 (red) are located in the most proximal 30% of CA1. All injections sites from mice in Figure 4 (blue) are located in the mid ~40%–60% of CA1.The most 30% of CA1 is the *Ten3*-high region of CA1 and the ~40%–60% of CA1 is the *Lphn2*-high region of CA1 as determined by *in situ* hybridization quantification on sagittal P8 brain sections (Figure S3).

**Table S1. Differential gene expression between Ten3-HIGH and Ten3-NONE cells across CA1, subiculum, and entorhinal cortex.** Results of differential expression analysis (Wilcoxon rank sum test) between Ten3-HIGH vs. Ten3-NONE cells in CA1, subiculum, or entorhinal cortex (individual tabs), with only genes more highly expressed in the Ten3-NONE group shown (avg_logFC < 0). To arrive at these lists, we analyzed each region separately, and defined Ten3-HIGH cells as those in the top $95^{th}$ percentile of Ten3 (Odz3) expression, and Ten3-NONE cells as those with no expression. In the columns of the table: ‘Gene’ is the name of the gene, ‘p_val’ is the p-value from the comparison, ‘avg_logFC’ is the average log fold change, ‘pct_Ten3-high’ is the fraction of Ten3-HIGH cells that expresses the gene, ‘pct_Ten3-none’ is the fraction of Ten3-NONE cells that express the gene, and ‘p_val_adj’ is the p-value with Bonferroni correction. We only considered genes that were expressed in at least 50% of cells in either of the comparison groups (min.pct = 0.5), and required that the difference in membership between the two groups (pct_Ten3-high – pct_Ten3-none) for considered genes to be >0.1 (min.diff.pct = 0.1).
